# Novel modification by L/F-tRNA-protein transferase (LFTR) generates a Leu/N-degron ligand in *Escherichia coli*

**DOI:** 10.1101/2021.07.26.453911

**Authors:** Ralf D. Ottofuelling, Robert L. Ninnis, Kaye N. Truscott, David A. Dougan

## Abstract

The N-degron pathways are a set of proteolytic systems that relate the half-life of a protein to its N-terminal (Nt) residue. In *Escherchia coli* the principal N-degron pathway is known as the Leu/N-degron pathway of which an Nt Leu is a key feature of the degron. Although the physiological role of the Leu/N-degron pathway is currently unclear, many of the components of the pathway are well defined. Proteins degraded by this pathway contain an Nt degradation signal (N-degron) composed of an Nt primary destabilizing (N_d1_) residue (Leu, Phe, Trp or Tyr) and an unstructured region which generally contains a hydrophobic element. Most N-degrons are generated from a pro-N-degron, either by endoproteolytic cleavage, or by enzymatic attachment of a N_d1_ residue (Leu or Phe) to the N-terminus of a protein (or protein fragment) by the enzyme Leu/Phe tRNA protein transferase (LFTR) in a non-ribosomal manner. Regardless of the mode of generation, all Leu/N-degrons are recognized by ClpS and delivered to the ClpAP protease for degradation. To date, only two physiological Leu/N-degron bearing substrates have been verified, one of which (PATase) is modified by LFTR. In this study, we have examined the substrate proteome of LFTR during stationary phase. From this analysis, we have identified several additional physiological Leu/N-degron ligands, including AldB, which is modified by a previously undescribed activity of LFTR. Importantly, the novel specificity of LFTR was confirmed *in vitro*, using a range of model proteins. Our data shows that processing of the Nt-Met of AldB generates a novel substrate for LFTR. Importantly, the LFTR-dependent modification of T_2_-AldB is essential for its turnover by ClpAPS, *in vitro*. To further examine the acceptor specificity of LFTR, we performed a systematic analysis using a series of peptide arrays. These data reveal that the identity of the second residue modulates substrate conjugation with positively charged residues being favored and negatively charged and aromatic residues being disfavored. Collectively, these findings extend our understanding of LFTR specificity and the Leu/N-degron pathway in *E. coli*.

## INTRODUCTION

Protein degradation is an essential cellular process that is responsible for the removal of unwanted or damaged proteins. Given the irreversible nature this process, the recognition of a protein substrate is generally tightly controlled, not only by the conditional exposure of a degron, but also by the regulated activation of distinct proteolytic machines that are responsible for recognition (and removal) of these proteins. In the bacterial cytosol, this process is performed by a handful of ATP-dependent machines, which are commonly referred to as AAA+ (<u>A</u>TPase <u>a</u>ssociated with a variety of cellular <u>a</u>ctivities) proteases (Striebel et al., 2009;Sauer and Baker, 2011;Gur et al., 2013;Alhuwaider and Dougan, 2017). These machines are generally composed of two components: an ATP-dependent unfoldase component belonging to the AAA+ superfamily (Neuwald et al., 1999;Ogura and Wilkinson, 2001), which is responsible for recognition and unfolding of the substrate and a specialized peptidase component responsible for destruction of the unfolded protein into short peptides. In a handful of cases, these machines also employ an additional component, commonly known as adaptor proteins, for the recognition of specific degrons (Dougan et al., 2002a;Kirstein et al., 2009;Mahmoud and Chien, 2018).

Degrons are generally short linear motifs (1 – 12 residues long) that serve as degradation signals. Given the main determinant of these signals is located at either the N- or C-terminus of a protein, they are commonly termed N- or C-degrons, respectively (Tobias et al., 1991;Tu et al., 1995;Keiler et al., 1996;Flynn et al., 2003;Erbse et al., 2006;Ninnis et al., 2009;Gao et al., 2019;Varshavsky, 2019;Timms and Koren, 2020). Although some degrons are constantly exposed and hence constitutively degraded, most degrons are generated conditionally, through the activation or exposure of a pro-degron (Hwang et al., 2010;Kim et al., 2014;Chen et al., 2017;Lucas and Ciulli, 2017;Dougan and Varshavsky, 2018;Varshavsky, 2019). The molecular components responsible for the generation, recognition and removal of a degron are defined by a degron pathway (Varshavsky, 2019;Timms and Koren, 2020). Currently, two N-degron pathways have been described in bacteria; the fMet/N-degron pathway for the co-translational removal of misfolded nascent polypeptides that retain their formyl group (Piatkov et al., 2015) and the Leu/N-degron pathway (Varshavsky, 2019), formerly the N-end rule pathway (Tobias et al., 1991), which is the canonical N-degron pathway in bacteria. Although the physiological role of the Leu/N-degron pathway in *E. coli* remains poorly understood, many of the molecular components are well defined. In the bacterial Leu/N-degron pathway, individual residues located at the N-terminus of a protein, can be considered either stabilizing or destabilizing (Tobias et al., 1991;Varshavsky, 2011). Similar to the Eukaryotic N-degron pathways, Nt destabilizing (N_d_) of the bacterial Leu/N-degron pathway are hierarchic (Mogk et al., 2007;Varshavsky, 2011;Dougan et al., 2012;Tasaki et al., 2012;Gibbs et al., 2014;Dissmeyer et al., 2018;Bouchnak and van Wijk, 2019;Varshavsky, 2019), composed of primary destabilizing (N_d1_) residues (Leu, Phe, Tyr and Trp) and secondary destabilizing (N_d2_) residues. While N_d1_ residues are recognized directly by ClpS, the N-recognin (N-terminal recognition component) of the pathway (Erbse et al., 2006;Wang et al., 2008b;Schuenemann et al., 2009), N_d2_ residues require specific modification to generate a destabilizing activity (i.e. attachment of a N_d1_ residue, e.g. Leu or Phe). To date, a total of five different N_d2_ residues have been identified in bacteria, three (Arg, Lys and in a single case, Met) in *E. coli* (Tobias et al., 1991;Shrader et al., 1993;Ninnis et al., 2009;Dougan et al., 2012) and four (Arg, Lys, Asp and Glu) in *Vibrio vulnificus* (Graciet et al., 2006). The modification of proteins bearing an N_d2_ residue is performed by two separate enzymes, the bacterial protein transferase (Bpt) is responsible for the Nt-leucylation of proteins bearing the N_d2_ residues (Asp or Glu) in *Vibrio vulnificus* (Graciet et al., 2006). While in *E. coli*, Leu/Phe-tRNA-protein transferase (L/F-transferase, here referred to as LFTR) is responsible for the conjugation of Leu or Phe to proteins bearing the N_d2_ residue (Arg, Lys or Met) (Tobias et al., 1991;Shrader et al., 1993;Ninnis et al., 2009;Schmidt et al., 2009). Based on the crystal structure, LFTR contains two pockets, one for the recognition of the acceptor (or substrate) and the other for recognition of the donor tRNA bearing the amino acid for conjugation to the substrate (Suto et al., 2006;Watanabe et al., 2007). While the donor specificity of LFTR *in vitro* includes Leu-tRNA, Phe-tRNA and to a lesser extent Met-tRNA (Kaji et al., 1963;Leibowitz and Soffer, 1970;Scarpulla et al., 1976), *in vivo* studies suggest that leucylation is the dominant type of conjugation (Shrader et al., 1993). Similarly, although the acceptor specificity of LFTR, originally defined using model substrates, was proposed to be restricted to the Nt amino acids Arg, Lys and to a lesser extent His (Soffer, 1973;Tobias et al., 1991), the identification of Putrescine aminotransferase (PATase, also known as PatA) as the first physiological substrate of LFTR showed that Met can also serve as an acceptor for this enzyme (Ninnis et al., 2009;Schmidt et al., 2009). This finding led to speculation that the acceptor specificity of LFTR may be broader than initially defined using model substrates (Tobias et al., 1991;Ninnis et al., 2009;Dougan et al., 2010). Therefore, in order to further investigate the acceptor specificity of LFTR we sought to identify the physiological substrates of this enzyme.

Here, we report the affinity isolation of ClpS-interacting proteins from an *E. coli* strain that lacks LFTR activity (Δ*aat*). Comparison of the ClpS-interacting proteins from this strain, with those isolated from Δ*clpA E. coli*, facilitated the identification of eight putative LFTR Leu/N-degron ligands. Three proteins (AldB, AccD and SufD) were verified, using specific antisera, as LFTR-dependent ClpS interacting proteins. The ClpAP-mediated *ex vivo* turnover of these proteins, was not only dependent on the presence of ClpS but also the activity of LFTR. Significantly, the presence of a non-ribosomal primary destabilizing residue (Leu) was confirmed by N-terminal sequencing of two ligands (AldB and AccD), and the ClpS-dependent turnover of these proteins was also verified using purified components. Unexpectedly, a primary destabilizing residue (Leu) was attached to an Nt Thr on AldB, which identifies MetAP is an integral component of the Leu/N-degron pathway. Furthermore, based on the identification of LT_2_AldB as a novel N-degron ligand, we systematically re-examined the *in vitro* specificity of LFTR using peptide-based arrays. Taken together, our data show that the recognition of N_d2_ residues by LFTR is broader than previously proposed and the specificity of LFTR is clearly influenced by residues immediately downstream of the N_d2_ residue of the acceptor protein. Finally, based on our identification of this novel ligand, we speculate that MetAP cleavage of other proteins bearing the Nt sequence, MTN may also be compatible with the generation of additional N-degron ligands (under different conditions). From this bioinformatic analysis, we identified 12 cytosolic proteins in *E. coli* with the Nt sequence MTN, six of which are (at least partially) cleaved by MetAP and one (BarA) is a putative ClpA-interacting protein (Butland et al., 2005;Rajagopala et al., 2014;Bienvenut et al., 2015) and hence may represent an additional Leu/N-degron substrate in *E. coli*.

## MATERIALS AND METHODS

### Strains, proteins, protein analysis and antibodies

*E. coli* knockout strains Δ*clpA* (JW0866) and Δ*aat* (JW0868), were grown at 37 °C for 26 h in LB media supplemented with 50 mg/ml kanamycin, as described in Ninnis et al., (2009). ClpA, ClpP and ClpS (wild-type and mutant) were expressed in *E. coli* and purified as described previously (Dougan et al., 2002b). LFTR was expressed and purified as described in Ninnis et al., (2009). Leu/N-degron substrates (and controls): SufD, K_24_SufD, LK_24_SufD AccD, K_16_AccD, LK_16_AccD, MT_2_AldB, T_2_AldB and LT_2_AldB, model GFP-fusion proteins: (LK_16_AccD_16-20_GFP, LK_16_AccD_16-24_GFP, LK_16_AccD_16-38_GFP, LK_16_AccD_16-55_GFP, MT_2_AldB_3-11_GFP, T_2_AldB_3-11_GFP and LT_2_AldB_3-11_GFP) and model LFTR substrates (and controls): R-PATase, T-PATase FL-PATase and FM-PATase were all generated using the Ub-fusion system (Catanzariti et al., 2004) and purified essentially as described in Ninnis et al., (2009). Coomassie-stained Leu/N-degron substrates were excised from 2D-SDS–PAGE gels and in-gel proteolytic digestion performed with either trypsin or GluC. Proteins were identified by MS/MS analysis as described in Ninnis et al., (2009). The N-terminal sequence of selected Leu/N-degron ligands was determined from a protein spot excised from a PVDF membrane, subjected to 5 – 7 cycles of automated Edman degradation, using an Applied Biosystems 494 Procise Protein sequencing system.

### In vitro transcription

The tRNA genes (*pheV* and *leu*Z) were amplified with specific primers that included a T7 promoter. Transcription of tRNA^pheV^ and tRNA^leuZ^ was performed with 20 U T7 RNA Polymerase (37 °C for 90 min) using the Riboprobe® *in vitro* Transcription System (Promega) essentially as described in the instructions manual. Following transcription, the sample (10 µl) was analyzed by gel electrophoresis using a 2 % (w/v) TAE-agarose gel to estimate the tRNA concentration.

### In vitro aminoacyl-transferase assay

Aminoacylation experiments were performed essentially as described (Ninnis et al., 2009), with minor modifications. For aminoacylation, the protein of interest (5 - 125 pmol) was incubated (37 °C for 8 min) in 25 µl reaction buffer (50 mM Tris-HCl pH 8.0, 100 mM KCl, 10 mM Mg(OAc)_2_, 1 mM DTT, 2 mM ATP) containing ∼1.0 µM of either tRNA^PheV^ or tRNA^LeuZ^, 8.75 µM [^14^C]-Phe/Leu (18.3 GBq/mmol (PerkinElmer), 38.5 U *E. coli* aminoacyl-tRNA synthetase (Sigma) and 0.18 µM Leucyl/Phenylalanyl-tRNA-protein transferase (LFTR). The reaction was stopped by the addition of sample buffer, then separated by 12.5 % Tris-glycine SDS-PAGE. Following separation, proteins were fixed (30 % (v/v) Methanol, 10 % (v/v) Acetic acid) in the gel for 30 min, then washed for 15 min in (30 % (v/v) Methanol, 2 % (v/v) Glycerol). After drying the gel (80 °C for 1.5 h) using a Model 583 Gel DRYER (Bio-Rad), it was exposed to a Phosphor Screen (GE Healthcare) for between 1 - 4 days and the protein signal visualized using a Typhoon Trio Variable Mode Imager (GE Healthcare).

To examine the binding specificity of LFTR, aminoacylation of peptides attached to a cellulose membrane was performed. Peptides, attached to a cellulose membrane through their C-terminus, were synthesized by spot synthesis (JPT Peptide Technologies). The N-terminal peptide sequences were derived from PATase, α-casein, β-galactosidase and AldB (see Supplementary Tables 3, 4 and 5 for peptide sequences of individual spots). The membrane was washed (three times with 1x PBS) prior to incubation with the reaction components (15 min at 37 °C, in a glass tube with gentle rolling). Prior to exposure of the membrane, the membrane was washed four times with 500 µl of 1x PBS. After air-drying the membrane was exposed to a Phosphor Screen (GE Healthcare) and the signal visualized using a Typhoon Trio Variable Mode Imager (GE Healthcare).

### In vitro degradation assay

Unless otherwise stated, *in vitro* degradation assays were routinely performed in 200 µl, using ClpAP Buffer (50 mM Tris-HCl pH 7.5, 300 mM NaCl, 20 mM MgAc, 10 % (v/v) Glycerol, 1 mM DTT) containing ClpA_6_P_14_ (200 nM) in the absence or presence of ClpS (1.2 µM). All reactions were pre-incubated (for 1 min at RT) with ATP (5 mM) to allow ClpAP complex formation, prior to the addition of the substrate. The reaction (performed at 37 °C) was initiated upon substrate addition (0.5 - 1 µM). To monitor the turnover of non-fluorescent protein substrates, samples were collected at various time-points (as indicated) and immediately mixed with SDS-PAGE loading buffer. Proteins were then separated by SDS-PAGE and visualized, either by staining with Coomassie Brilliant Blue or by immunodecoration with specific antisera following transfer to a PVDF membrane. An ATP-regeneration system (4 mM Phosphoenolpyruvate (Sigma) and 20 µg/ml pyruvate kinase (Sigma)) was included in reactions lasting longer than 60 min. To monitor the turnover of fluorescent substrates (e.g. GFP-tagged protein substrates) GFP fluorescence (excitation wavelength = 400 nm and emission wavelength = 510 nm) was monitored for the indicated times using a Spectramax M5e plate reader (Molecular Device Inc.), essentially as described (Dougan et al., 2002b).

### Purification of ClpS interacting proteins

To study N-degron binding, *in vitro* “pull-down” experiments were performed as described previously (Geissler et al., 2002;Ninnis et al., 2009). Briefly, settled NiNTA-agarose beads (QIAGEN) were equilibrated in Buffer A (50 mM Tris-HCl pH 8.0, 300 mM NaCl, 5 mM Imidazole). The bait protein (wild type or mutant His_6_-ClpS or His_10_-ClpS) was immobilized to the equilibrated beads (15 min end-over-end mixing at 4 °C) at a ratio of 2 µg of bait protein per 1 µl of settled beads. The beads were then washed (3 × 10 min) with 3 bed volumes (BV) of Buffer B (50 mM Tris-HCl pH 8.0, 300 mM NaCl, 20 mM Imidazole) followed by a single wash (10 min at 4 °C) with end-over-end mixing using 3 BV of Buffer C (20 mM HEPES-KOH pH 7.5, 100 mM KOAc, 10 mM Mg(OAc)_2_, 10 % (v/v) Glycerol, 10 mM Imidazole, 0.5 % (v/v) Triton X-100). For N-degron binding studies using purified substrate proteins, NiNTA-agarose beads containing immobilized His_6_- or His_10_-ClpS were incubated (30 min at 4 °C, with end-over-end mixing) with an equimolar amount of the prey protein. To isolate novel ClpS-interacting proteins, ∼1 g of soluble *E. coli* cell lysate (in Buffer C supplemented with a cocktail of protease inhibitors (cOmplete, EDTA-free (Roche)), was incubated (30 min at 4 °C) with immobilized ClpS (∼1 mg) with end-over-end mixing. Unbound proteins were removed by centrifugation (300 g for 5 min at 4 °C) and the slurry containing bound proteins transferred to a 1 ml MoBiTec column (Molecular Biotechnology). The slurry was washed with 40 BV of Buffer D (Buffer C containing 0.25 % (v/v) Triton X-100), residual buffer was removed by centrifugation (300 g for 1 min at 4 °C). Finally, ClpS-interacting proteins were eluted by centrifugation (300 g for 1 min at 4 °C) with 1 BV of FR-dipeptide (1 mg/ml) in Buffer C (without Triton X-100). Eluted proteins were analyzed by SDS-PAGE, immunoblotting or 2D-PAGE.

### 2D-PAGE

ClpS-interacting proteins (max. 250 µg), recovered by FR dipeptide elution, from an *E. coli* cell lysate were precipitated with 4 volumes of cold acetone and resuspended in 150 µl rehydration solution (8 M Urea, 2 % (w/v) CHAPS, 0.5 % (v/v) IPG buffer 4 - 7 or 3 – 10 (Pharmacia), 20 mM DTT, 0.002 % (w/v) Bromophenol blue). Rehydrated protein samples were separated according to their isoelectric point on an Immobiline® DryStrip gel (13 cm, linear pH 4-7 or 3-10 gradient strip (Pharmacia)) using an Ettan IPGphor II Manifold with cup loading. The samples were loaded towards the anode end of the rehydrated DryStrip gel (in rehydration solution for 10-20 h) and the proteins focused using the following conditions: 100 V for 0.5 h, from 100 V to 500 V over 2 h, from 500 V to 1,000 V over 1 h, from 1,000 to 8,000 V over 3.5 h and 8,000 V for 1 h at 20 °C. The DryStrip gel was then equilibrated with gentle rocking, first in Buffer A (50 mM Tris-HCl pH 8.8, 6 M Urea, 30 % (v/v) Glycerol, 1 % (w/v) DTT and 2 % (w/v) SDS) then in Buffer B (50 mM Tris-HCl pH 8.8, 6 M Urea, 30 % (v/v) Glycerol, 135 mM Iodoacetamide and 2 % (w/v) SDS), each for 15 min. The equilibrated DryStrip gel was then placed on top of a 4-16 % Tris-Tricine gel and the proteins separated, in the second dimension, by SDS-PAGE and visualized by Coomassie Brilliant Blue staining or immunodecoration after being transferred to PVDF.

## RESULTS

### Deletion of *aat* (encoding LFTR) inhibits docking of specific N-degron ligands to ClpS

Although there have been significant advances defining the physiological role of the Leu/N-degron pathway in Salmonella (Yeom et al., 2017;Gao et al., 2019;Yeom and Groisman, 2019), our current understanding of this pathway in *E. coli* is largely derived from *in vitro* studies using model substrates (Tobias et al., 1991;Shrader et al., 1993;Erbse et al., 2006;Wang et al., 2008b;Kress et al., 2009;Schuenemann et al., 2009;Roman-Hernandez et al., 2011;Varshavsky, 2011;Rivera-Rivera et al., 2014). As a consequence, the physiological substrates of ClpS are largely unknown and the biological function of the pathway is currently unclear (Ninnis et al., 2009;Schmidt et al., 2009;Dougan et al., 2010;Humbard et al., 2013). Previously, we developed an affinity method to isolate and identify physiological Leu/N-degron substrates from *E. coli* (Ninnis et al., 2009). To help determine, which of the previously identified ligands may be *bona fide* N-degron substrates of the ClpAPS machinery, we generated antibodies to a selection of ligands and monitored their ClpS-dependent turnover by ClpAP *ex vivo* (**Figure 1A** and **B**). From these experiments we identified SufD (of the SufC/D complex), AccD (of the AccA/D complex) and AldB as putative Leu/N-degron substrates. To determine which of the above putative Leu/N-degron substrates are modified by LFTR we isolated Leu/N-degron ligands from Δ*clpA* cells and compared them with the Leu/N-degron ligands from a mutant *E. coli* strain (Δ*aat*) which lacks LFTR (**Figure 1C** and Supplementary Figure 1). Initially we analyzed the ClpS-interacting proteins by SDS-PAGE (**Figure 1C**). Consistent with our previous analysis, ∼ 30 different proteins were eluted (using the FR dipeptide) from the wild type ClpS column (**Figure 1C**, compare lanes 4 and 5). This included two highly abundant proteins (Dps at ∼ 17 kDa and PATase at ∼ 50 kDa), previously identified as natural substrates of the *E. coli* Leu/N-degron pathway (Ninnis et al., 2009;Schmidt et al., 2009). Next, we used 2D-PAGE to compare the ClpS-interactome isolated from Δ*clpA* and Δ*aat* cells. As a control, we monitored the recovery of PATase, a confirmed LFTR-dependent substrate (Ninnis et al., 2009;Schmidt et al., 2009;Humbard et al., 2013). As expected, and consistent with our previous findings, PATase was absent from the dipeptide eluted fraction derived from Δ*aat* cells (**Figure 1C**, lane 6 and Supplementary Figure 1B, Spot 3). These data validate the strains used for the isolation of Leu/N-degron substrates and our approach to identify Leu/N-degron substrates that are modified by LFTR. Notably, more than half of the prominent protein spots were essentially unchanged in the two elution profiles (Supplementary Figure 1C, black Spots 9 - 17) suggesting that, under these conditions, the majority of Leu/N-degron ligands are not modified by LFTR. Nevertheless, using this approach we were able to identify nine prominent protein spots (recovered from Δ*clpA* cells) that were absent from the dipeptide eluted fraction derived from Δ*aat* cells (Supplementary Figure 1C, black spots, red numbers 1 – 8), suggesting that several N-degron ligands are modified by LFTR *in vivo*. To determine which proteins were modified by LFTR and at which residue this modification occurred, we identified the proteins recovered from Δ*clpA* by Mass Spectrometry and determined the N-terminal sequence of the most prominent spots (Supplementary Figure 1C, dotted red circles, see Supplementary Table 1). From these data, we identified eight LFTR-dependent ligands (see Supplementary Table 1), two of which (AccA and SufC) were excluded as *bona fide* N-degron substrates, based on the absence of an N_d1_ residue (i.e. the Nt residue of AccA, recovered from the pull-down, was Ser2) or by the lack of ClpAPS-mediated turnover (i.e. although both SufC and SufD were both recovered by pull-down in an LFTR-dependent manner, only SufD was degraded by ClpAPS, *ex vivo*) (see **Figure 1B** and **C**). Of the remaining six LFTR-dependent ligands, the N-terminal sequence of three (AccD, AldB and PATase) was experimentally determined (Supplementary Table 1) while the N-terminus of two other proteins (SufD and RsgA) was proposed, based either on the apparent MW of the recovered ligand or published evidence (Supplementary Table 1). Unfortunately, we were unable to identify the putative N-degron within ClpB.

**Figure 1.**
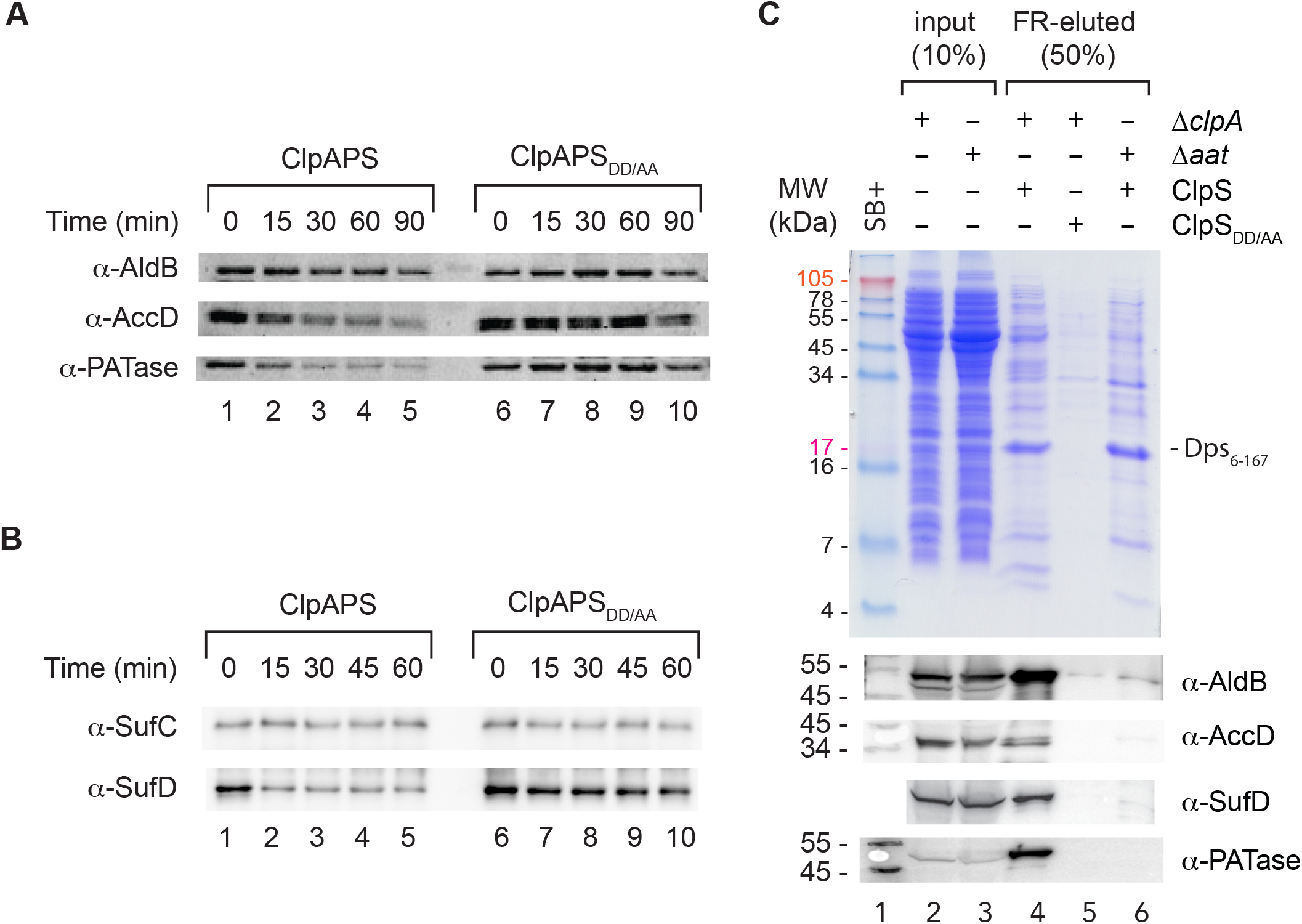
Identification of putative LFTR-dependent N-degron substrates from *E. coli*. **(A)** The *ex vivo* turnover of selected *E. coli* N-degrons; AldB, AccD and PATase (as a positive control) was monitored using specific antisera, in the presence of ClpAPS (lanes 1 – 5) or ClpAPS_DD/AA_ (lanes 6 – 10). **(B)** The *ex vivo* turnover of putative *E. coli* N-degrons (SufC and SufD) was monitored using specific antisera, in the presence of ClpAPS (lanes 1 – 5) or ClpAPS_DD/AA_ (lanes 6 – 10). **(C)** *E. coli* proteins from either a Δ*clpA* (lane 2) or Δ*aat* (lane 3) *E. coli* cell lysate, were applied to Ni-NTA agarose beads containing immobilized wild type (lanes 4 and 6) or mutant (lane 5) ClpS. N-degron proteins were specifically eluted from wild type ClpS (lanes 4 and 6) and not from the immobilized mutant, ClpS_DD/AA_ (lane 5). LFTR-dependent N-degrons (AldB, AccD and SufD) were only recovered in the FR-eluted fraction from Δ*clpA* cells (lane 4) and not in the FR-eluted fraction from Δ*aat* cells (lane 6). Proteins were separated by Tricine SDS–PAGE.

In the case of SufD, although its interaction with ClpS was dependent on LFTR activity and its *ex vivo* turnover was ClpAPS-dependent (**Figure 1B**) we were unable to determine the identity of its N-terminal residue. Therefore, based on the apparent MW of SufD (recovered from the pull-down) we identified a potential processing site and speculated that SufD was processed (by an unknown peptidase) to reveal Lys24 at the N-terminus (i.e. K_24_-SufD), to which an N_d1_ residue is attached. Consistent with this proposal, recombinant LK_24_-SufD co-migrated with processed SufD recovered from the pulldown (Supplementary Figure 2A) and was rapidly degraded *in vitro* by ClpAPS (Supplementary Figure 2B, filled circles and Supplementary Figure 2C). Interestingly, although the post-translational modification of SufD was not essential for its turnover, the type of modification did control the rate of SufD turnover *in vitro*. For instance, in the absence of endoproteolytic processing, the ClpAP-mediated turnover of SufD was very slow (Supplementary Figure 2B, open triangles). However, following removal of the N-terminal segment, the rate of SufD turnover (i.e. K_24_-SufD) was enhanced, by ∼ 2.5-fold. Notably, the turnover of both SufD and K_24_-SufD was completely inhibited in the presence of ClpS (Supplementary Figure 2B, filled triangles and filled squares), while in contrast, LK_24_-SufD was rapidly degraded in the presence of ClpS. Indeed, in the presence of ClpS the ClpAP-mediated turnover of SufD was increased ∼ 6-fold by its processing and modification (Supplementary Figure 2B, compare open triangles and filled circles). Taken together these *in vitro* data could suggest that processing of SufD, through activation of the Leu/N-degron pathway, is a potential mechanism to fine-tune the rate of SufD turnover in the presence of ClpS and hence control the cellular levels of SufD. However, the physiological conditions that might trigger SufD processing and its conversion into a putative N-degron substrate currently remain unknown.

Next, we examine the *in vitro* turnover of LK_16_-AccD relative to full length AccD (Supplementary Figure 3). Similar to SufD, full length AccD appears to contain a weak ClpA-recognition motif as it is slowly degraded (t_½_ > 4 h) by ClpAP in the absence of ClpS (Supplementary Figure 3A, open squares). Interestingly, this recognition motif appears to be located within this first 15 residues of AccD, as removal of these residues prevents its turnover by ClpAP (Supplementary Figure 3A, open circles). In contrast, attachment of an N_d1_ residue to processed AccD (LK_16_AccD) generates a classic Leu/N-degron substrate, which is specifically and rapidly (t_½_ ∼ 8 min) degraded by ClpAP in the presence of ClpS (Supplementary Figure 3A, filled circles). Given the N-terminal region of AccD (residues 23-50) contains a stable C4-type Zn-finger domain, we were interested to understand how processed AccD is delivered to ClpAP. To do so, we examined the sequence and structure of the AccD C4-type Zn-finger domain. From this analysis we identified a hydrophobic patch on the surface of AccD, composed of two discontinuous hydrophobic sequences (Supplementary Figure 3C). To examine the potential involvement of these sequence elements in substrate delivery to ClpA(P), we generated a series of GFP-fusion proteins which contained N-terminal segments (of different lengths) derived from LK_16_AccD (see Supplementary Figure 3B). The shortest segment contained only 5 residues (and lacked the first hydrophobic element). The next construct contained 4 additional residues (9 in total and included the first hydrophobic sequence, VW). Finally, the longest construct included the entire C4-type Zn-finger domain (and both hydrophobic sequences) while the last construct was intermediate in size but still included both hydrophobic elements. Interestingly, although the shortest construct lacked both hydrophobic elements, some turnover by ClpAPS was still observed (Supplementary Figure 3C, open blue circles), suggesting that a hydrophobic element is not essential for delivery of all N-degron substrates to ClpA(P). An alternate explanation for this result, may be that a dihydrophobic element (LF) near the N-terminus of GFP can act as surrogate for delivery to ClpAP. Nevertheless, the rate of turnover was dramatically enhanced when the first hydrophobic element (VW) was included in the sequence (Supplementary Figure 3C, open red circles), and the delivery was further improved when both hydrophobic elements were included in the GFP-fusion protein. Collectively, these data suggest that a linker sequence with at least nine residues downstream of the N_d1_ residue is required for efficient delivery to ClpA. Importantly, these data are consistent with previous findings from several groups, showing that the length of the linker region plays a critical role in substrate handoff to ClpAP (Erbse et al., 2006;Wang et al., 2008a;Ninnis et al., 2009).

In summary, consistent with the current dogma for the generation of Leu/N-degron substrates in bacteria, both AccD and SufD are generated from a pro-N-degron, via an unknown endopeptidase, which reveals a classic N-terminal secondary destabilizing (N_d2_) residue – Lys – (i.e. K_16_ in AccD and K_24_ in SufD), to which a primary destabilizing (N_d1_) residue (L or F) is then attached by LFTR to generate an N-degron ligand (i.e. LK_16_-AccD or LK_24_-SufD). In contrast to AccD and SufD, the N-degron of AldB is generated by an exopeptidase to remove a single residue, the initiating Met, which exposes Thr2 at the new N-terminus. This activity is consistent with processing by MetAP, at what has been termed a “twilight” residue (Frottin et al., 2006;Bienvenut et al., 2015;Yang et al., 2019). Hence, it appears that MetAP processing of AldB generates a non-canonical substrate (T_2_-AldB) for attachment of a primary destabilizing residue (L) to produce an N-degron ligand (LT_2_-AldB). Unexpectedly, the conjugation (of Leu or Phe to T_2_-AldB) was dependent on the activity of LFTR *in vivo*. Therefore, in order to confirm the potential of this conjugation with respect to N-degron degradation, we first generated a series of recombinant proteins (using the Ub-fusion system, to ensure the identity of the N-terminal residue) and monitored the ClpAP-dependent turnover of these proteins in the absence or presence of ClpS (**Figure 2**). Consistent with our identification of LT_2_-AldB as both a ClpS-ligand and ClpS-dependent substrate of ClpAP *ex vivo*, recombinant LT_2_-AldB was only degraded by ClpAP in the presence of ClpS (**Figure 2A**, lower panel lanes 8 – 14). In contrast, both unprocessed AldB (MT_2_-AldB) and MetAP-processed AldB (T_2_-AldB) were stable both in the presence and absence of ClpS (**Figure 2A**, upper and middle panels, respectively). Interestingly, similar to AccD and SufD, a compelling hydrophobic element was also absent from the N-terminal region of AldB, therefore to examine if a hydrophobic element was essential for substrate delivery to ClpA, we fused the first 11 residues of X-AldB (where X refers to either MT, T or LT) to GFP (**Figure 2B**). Consistent with the turnover of authentic LT_2_-AldB, LT_2_AldB_3-11_GFP was the only GFP-fusion protein to be degraded by ClpAPS (**Figure 2B**, black open circles). Collectively these data confirm LT_2_-AldB as an N-degron substrate and demonstrate that delivery of this N-degron substrate to ClpA(P) can occur in the absence of a “strong” hydrophobic element within the linker region.

**Figure 2.**
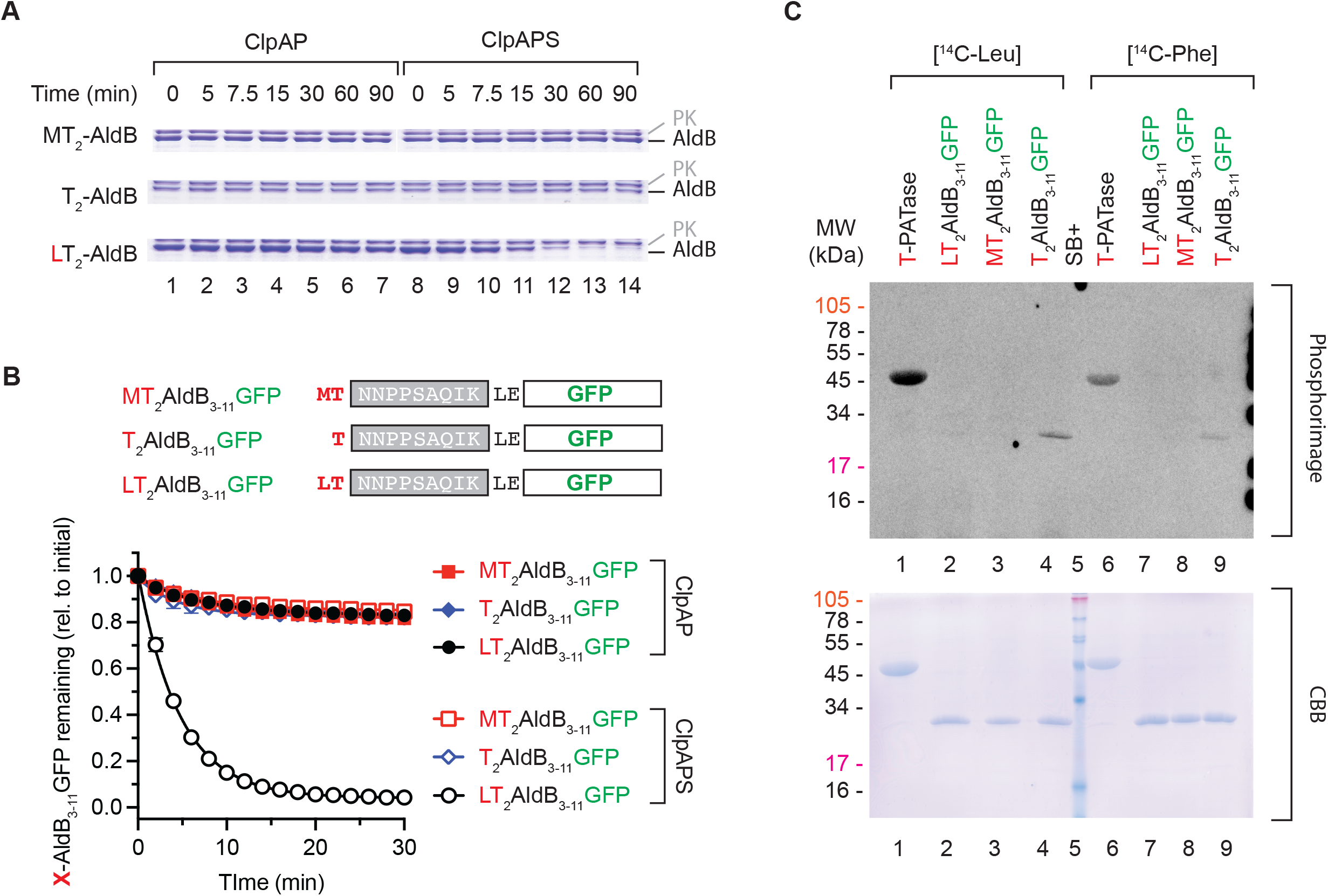
The LFTR-dependent leucylation of T_2_-AldB generates a ClpS-dependent substrate for ClpAP, *in vitro*. **(A)** The *in vitro* turnover of AldB is dependent on the presence of an N_d1_ (Leu). The ClpAP-mediated turnover of recombinant X-AldB was monitored *in vitro*, in the absence (lanes 1 – 7) or presence (lanes 8 – 14) of ClpS. Only LT_2_-AldB (lower panel) was degraded by ClpAPS, neither MT_2_-AldB (upper panel) nor T_2_-AldB (middle panel) were degraded by ClpAP or ClpAPS. **(B)** Schematic representation of the GFP fusions (upper panel). The turnover of MT_2_-AldB_3-11_GFP (red squares), T_2_-AldB_3-11_GFP (blue diamonds) and LT_2_-AldB_3-11_GFP (black circles) was monitored in the absence (filled symbols) or presence (open symbols) of ClpS. Protein turnover was monitored by the loss of GFP fluorescence (λ_ex_ = 400 nm and λ_em_ = 510 nm). **(C)** The LFTR-dependent modification of T-PATase (lanes 1 and 6), LT_2_-AldB_3-11_GFP (lanes 2 and 7), MT_2_-AldB_3-11_GFP (lanes 3 and 8) and T_2_-AldB_3-11_GFP (lanes 4 and 9) was monitored in the presence of either [^14^C]-Leu (lanes 1 – 4) or [^14^C]-Phe (lanes 6 – 9). See blue MW markers are indicated.

Having confirmed that LT_2_-AldB is a potential N-degron substrate, we examined the ability of Thr to act as a secondary destabilizing residue for LFTR. Initially as a control, we tested the activity of LFTR using two artificial model substrates (GFP and PATase) bearing a classic secondary destabilizing residue (R) at the N-terminus (i.e. R-βgal_2-11_-GFP and R-PATase). As expected, [^14^C]-Leu was attached to R-βgal_2-11_-GFP (Supplementary Figure 4A, lane 3) and R-PATase (Supplementary Figure 4B, lane 3). Next, we used PATase as a model protein to examine the ability of N-terminal Thr to act as an acceptor for LFTR. Consistent with our identification of LT_2_-AldB as a N-degron ligand recovered from Δ*clpA* cells, T-PATase served as an acceptor for the LFTR-dependent attachment of radiolabeled Leu (Supplementary Figure 4A, lane 2). We then examined the ability of LFTR to catalyze the attachment of [^14^C]-Phe to a selection of X-PATase fusion proteins (Supplementary Figure 4B). Consistent with the conjugation of [^14^C]-Leu, [^14^C]-Phe was also attached to both T- and R-PATase (albeit to a reduced level than [^14^C]-Leu), but not to FM-PATase (Supplementary Figure 4B). Finally, we examined the conjugation of [^14^C]-Leu (or [^14^C]-Phe) to recombinant T_2_-AldB, however despite our efforts we were unable to reconstitute this system *in vitro*. As a result, we speculated that the lack of conjugation to T_2_-AldB may be due to restricted accessibility of the N-terminus of AldB to LFTR, which could serve as a mechanism to regulate its conjugation *in vivo*. To overcome this potential constraint, we tested the ability of an Nt Thr (in the context of the native AldB sequence) to act as substrate for LFTR, using the X-AldB_3-11_GFP fusion protein. Importantly, both radiolabeled amino acids ([^14^C]-Leu and [^14^C]-Phe) were conjugated to T-AldB_3-11_GFP (**Figure 2C**, lanes 4 and 9), while in contrast neither LT-AldB_3-11_GFP (**Figure 2C**, lanes 2 and 7) nor MT-AldB_3-11_GFP (**Figure 2C**, lanes 3 and 8) served as an acceptor for the conjugation of either amino acid. Collectively, these data confirm that an N-terminal Thr residue can serve as an acceptor for LFTR and suggests that the acceptor specificity of LFTR is broader than originally proposed (Shrader et al., 1993). Interestingly, both non-canonical LFTR substrates (i.e. PATase and AldB) shared the same downstream residue (N). As such, we speculated that (a) the identity of the second residue (adjacent to the N_d_) might contribute to LFTR specificity/activity and (b) more specifically, substrates/proteins bearing a non-canonical Nt-residue might exhibit a restricted preference for specific residues in position 2.

Therefore, in order to gain a more complete understanding of LFTR acceptor specificity, we examined the LFTR-dependent conjugation of [^14^C]-Leu to several libraries of cellulose bound 11-mer peptides. Initially, as a control, we examined the conjugation of [^14^C]-Leu to 11-mer peptides derived from the well-established model peptide substrate, casein fragment 90 – 95 (which includes an Nt Arg) and compared the conjugation to a series of related peptides in which the Nt residue was exchanged for each of the remaining 19 amino acids. All peptides, derived from the casein fragment 90 – 95 also contained the sequence AGSAG at positions 7–11. Initially we examined the specificity using a peptide library arranged in functional groups (Supplementary Figure 5A). As expected, the peptides bearing an Nt basic residue (Arg or Lys) served as an acceptor for LFTR (Supplementary Figure 5A, spots A-35 and A-36). To ensure the observed conjugation specificity was not due to an uneven distribution of reaction components over the membrane, we altered the arrangement of immobilized peptides on the cellulose membrane (Supplementary Figure 5C, spots 44 and 53). Consistent with Supplementary Figure 5A, the activity of LFTR was unchanged by the peptide position on the cellulose membrane (Supplementary Figure 5C). Importantly, even following prolonged exposure of the membrane(s), incorporation of [^14^C]-Leu in “negative” peptides spots was not observed. This suggests that low levels of conjugation are likely to represent actual LFTR-mediated conjugation. Next, we examined the specificity of the residue downstream of N_d2_, while maintaining Arg at the N-terminus of the casein peptide (**Figure 3A** and Supplementary Figure 6) or a peptide derived from the model protein β-gal (Supplementary Figure 7). Consistent with the idea that the residue at position 2 of the substrate, modulates LFTR specificity, the conjugation of [^14^C]-Leu varied depending on the identity of this residue. In fact, based on the relative activity of LFTR, the residue downstream of N_d2_ (at position 2) could be broadly categorized into three groups (favored, accepted and disfavored residues). While basic residues (Arg and Lys) were the most favored position 2 residue, polar residues (i.e. Ser, Thr, Gln and Asn), Gly, Pro and His were also the accepted. In contrast to these residues, small hydrophobic residues (i.e. Ala, Leu, Ile and Met) were only weakly accepted at position 2 of the substrate, with relative conjugation rates of ∼ 50% (Supplementary Figure 8). In contrast to these accepted residues, the remaining residues (i.e. acidic, aromatic and Cys) were all disfavored, with acidic residues the most strongly disfavored residue at position 2. Overall, these changes in the level of conjugation (to two sets of different 11-mer peptides bearing an Nt Arg) clearly demonstrate that (for substrates bearing a classic N_d2_ residue, Arg) the identity of the downstream residue does contribute to LFTR activity. These data are also consistent with the idea that the identity of this residue also plays an important role in modulating the specificity of LFTR in the recognition of substrate proteins bearing a non-canonical N_d2_ residue (i.e. Met or Thr). Indeed, the influence of the residue downstream of on LFTR specificity, for protein substrates bearing non-canonical N_d2_ residues, may be greater than that for canonical N_d2_ residues (i.e. Arg and Lys). Furthermore, although His was a permissive residue at position 2 in the context of an Nt Arg, we noted a small but specific difference in the conjugation of [^14^C]-Leu to RH-casein in comparison to RH-β-gal, which was not observed with any other amino acid at position 2. Although we cannot exclude that this small difference may be due to the limited sample size of these single use, peptide array experiments, there is remarkable consistency across the remaining 19 amino acids. Therefore, we propose that the residue(s) downstream of position 2 (i.e. position 3 and 4), may also make a minor contribution to LFTR specificity that may be particularly important for LFTR-substrates bearing a non-canonical N_d2_ residue.

**Figure 3.**
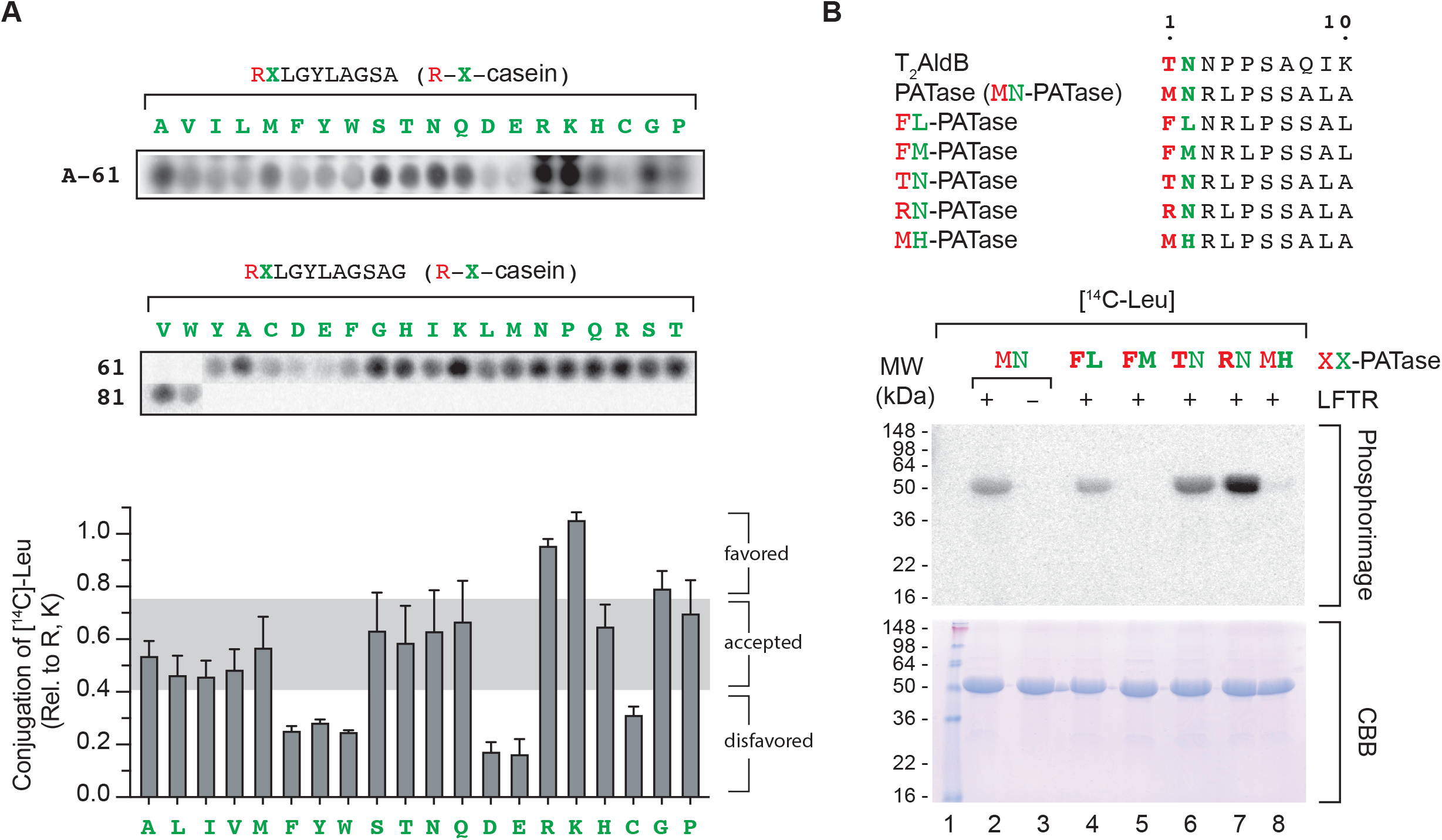
The specificity of LFTR is influenced by the identity of the residue in position 2. **(A, upper panel)** [^14^C]-Leu phosphorimages of two 11-mer peptide libraries with LFTR highlighting R-X-casein peptide spots (for the full phosphorimage of each library, see Supplementary Figure 6). Peptide sequences for R-X-casein peptides are indicated below each peptide library panel. For details of all other peptide sequences see Supplementary Tables 2 and 3. **(A, lower panel)** Conjugation of [^14^C]-Leu to R-X-casein, relative to the average conjugation of R-b-casein, where b = R, K. Relative conjugation activity was determined from two independent experiments and is separated into three broad categories (favored (>75%), accepted (40 – 75%, grey panel) and disfavored (< 40%)). **(B)** The Nt leucylation of XX-PATase *in vitro* is dependent on the identity of the first two residues. Nt sequences of XX-PATase mutants used in the assays (upper panel). Recombinant XX-PATase was separated by 12.5% SDS-PAGE and stained with Coomassie Brilliant Blue (CBB, lower panel). Following drying of the stained, polyacrylamide gel, the [^14^C]-Leu radiolabeled proteins were detected by phosphor image analysis using a Typhoon Trio Variable Mode Imager (panel). As a control MN-PATase was incubated in the presence (lane 2) or absence (lane 3) of LFTR. All other XX-PATase variants were incubated in the presence of LFTR. FL-PATase (lane 4), FM-PATase (lane 5) TN-PATase (lane 6) RN-PATase (lane 7) and MH-PATase (lane 9). See blue + MW markers (lane 1).

Therefore, to further examine the role of His (at position 2) in a substrate bearing a non-canonical N_d2_, we monitored the *in vitro* conjugation of [^14^C]-Leu to a selection of recombinant PATase mutant proteins, in which either the N-terminal or the second residue of the protein was altered (**Figure 3B upper panel**). As expected, the conjugation of [^14^C]-Leu was strongest to the positive control acceptor protein (i.e. RN-PATase), which contains a canonical N_d2_ residue followed by an “accepted” position 2 residue (**Figure 3B lower panel**, lane 7). Similarly, the conjugation of [^14^C]-Leu (or [^14^C]-Phe) was not observed to the negative control acceptor protein (i.e. FM-PATase), which bears a bulky hydrophobic residue at both the N-terminus and position 2 (**Figure 3B lower panel**, lane 5 and Supplementary Figure 9). In contrast, weak conjugation of [^14^C]-Leu (but none of [^14^C]-Phe) was observed for the acceptor FL-PATase (**Figure 3B lower panel**, lane 4). Although these data are somewhat surprising, they are consistent with the poly-leucylation (conjugation of multiple Leu residues) observed for M-PATase, LM-PATase and LLM-PATase both *in vitro* and *in vivo* (Ninnis et al., 2009;Humbard et al., 2013) which demonstrates that although the acceptor pocket of LFTR is able to accommodate two small hydrophobic residues, it is unable to accommodate two large hydrophobic residues. Most importantly, and consistent with a moderating role for position 2 of the acceptor, the conjugation of [^14^C]-Leu to PATase, was completely inhibited by its replacement with His (i.e. MH_2_-PATase) (**Figure 3B lower panel**, compare lanes 2 and 8). A similar profile was also observed for the conjugation of [^14^C]-Phe to the same protein substrates, albeit with a reduced activity (Supplementary Figure 9). Collectively, these data confirm that LFTR-substrates bearing a non-canonical N_d2_ residues (i.e. M or T, L or F) are influenced by the identity of the downstream residue, and more specifically that Asn appears to be preferred over His, in the context of a substrate bearing an Nt Met.

## DISCUSSION

In this study, we used ClpS-affinity chromatography to isolate Leu/N-degron ligands from Δ*clpA* and Δ*aat E. coli* cells in stationary phase. Comparison of the dipeptide eluted proteins recovered from Δ*clpA* cells and not from Δ*aat* cells identified eight ligands that are dependent on LFTR activity for their interaction with ClpS (**Figure 1**). Of these eight LFTR-dependent Leu/N-degron ligands, four (AldB, AccD, SufD and PATase) are degraded by ClpAPS *ex vivo*, three of which (AldB, AccD and PATase) we confirmed by N-terminal sequencing to contain a non-ribosomal N_d1_ residue (i.e. Leu). In addition to the above proteins, ClpB and EngS were also identified as potential LFTR-dependent substrates, however as we were unable to determine the N-terminal sequence of these protein or monitor their turnover *ex vivo*, these proteins remain unverified Leu/N-degron ligands. In contrast to the above proteins, AccA and SufC are identified as passenger ligands that co-purified with genuine Leu/N-degron ligands (AccD and SufD, respectively). Of the confirmed Leu/N-degron ligands, PATase was previously identified as a LFTR-dependent substrate in which the initiating Met was shown to serve as an N_d2_ residue (Ninnis et al., 2009;Schmidt et al., 2009;Humbard et al., 2013). The three remaining ligands are processed prior to their conjugation by LFTR. For AccD and SufD the processing involves removal of a short N-terminal segment, via an unidentified endopeptidase. While in the case of AldB, removal of the initiating Met by the exopeptidase MetAP is sufficient to generate an LFTR substrate. Unexpectedly, N-terminal sequencing of AldB (eluted from the ClpS affinity column) revealed that the primary destabilizing residue (Leu) is post-translationally attached to Thr2 of AldB. Hence, these data demonstrate that AldB is a novel LFTR-dependent Leu/N-degron ligand, and show that, like the eukaryotic N-degron pathways, MetAP plays a direct role in the bacterial Leu/N-degron pathway (Varshavsky, 2011;Nguyen et al., 2019;Varshavsky, 2019). Importantly, although the LFTR-dependent modification of recombinant T_2_-AldB was not confirmed, the leucylation of two model proteins (bearing an Nt Thr): i.e. T-PATase and a T_2_-AldB_3-11_GFP (**Figure 2**), was observed *in vitro*. One explanation for this apparent incongruity is that the N-terminus of recombinant T_2_-AldB is inaccessible to LFTR *in vitro* and the modification of T_2_-AldB *in vivo* is conditional upon exposure of its N-terminus. Importantly, consistent with the identification of LT_2_-AldB as an LFTR-dependent Leu/N-degron ligand, LT_2_-AldB is rapidly degraded by ClpAPS *in vitro* (**Figure 2**) although we have yet to establish the condition for AldB turnover *in vivo*. Taken together, our data suggest that T_2_-AldB is a conditional LFTR-dependent substrate, the modification of which is dependent on exposure of the N-terminus and the identity of the residue in position 2. This conditional recognition is somewhat reminiscent of the Ac/N-degron pathway in mammals, in which Ac/N-degrons only become exposed (and hence degraded) for instance, in the absence of a partner protein (Hwang et al., 2010;Shemorry et al., 2013;Nguyen et al., 2018). Based on our findings, we propose a model for the conditional modification and degradation of *E. coli* AldB (Supplementary Figure 10). In this model, Nt Met excision (by MetAP) is a crucial step in preparing T_2_-AldB for its conditional modification by LFTR. This modification of T_2_-AldB generates a Leu/N-degron ligand (LT_2_-AldB) which is recognized by ClpS and degraded by ClpAP *in vitro*. Therefore our findings suggest, that in addition to the basic residues (Arg and Lys) (Tobias et al., 1991) and the initiating Met of PATase (Ninnis et al., 2009), Nt Thr (of AldB) can also act as a N_d2_ residue for conjugation by LFTR (during stationary phase).

What is the function of *E. coli* AldB and why is T_2_-AldB modified by LFTR? AldB belongs to a group of enzymes (Aldehyde dehydrogenases), that catalyze the oxidation of aldehydes to carboxylic acids. Currently, the function of AldB in *E. coli* is unclear, however its expression has been linked to persister cell formation (Kawai et al., 2018) and a short-term adaptation response to glucose limiting conditions (Franchini and Egli, 2006). The expression of AldB is also upregulated in response to ethanol stress and upon entry into stationary phase and as such has been proposed to detoxify alcohols and aldehydes that accumulate during stationary phase (Xu and Johnson, 1995;Ho and Weiner, 2005). Interestingly, YiaY (encoded by the gene upstream of *aldB*) is a putative alcohol dehydrogenase, which together with AldB, contributes to sequential enzymatic steps in the oxidation of ethanol to acetate, via acetaldehyde. Therefore, one possibility is that following recovery from ethanol stress or on exit from stationary phase the cellular levels of AldB (and YiaY) are controlled by the Leu/N-degron pathway. Intriguingly, YiaY is also known to exhibit Threonine dehydrogenase (TDH) activity (Ma et al., 2014), and TDH activity in *E. coli* was previously proposed to be regulated by LFTR (Newman et al., 1976), Despite this, a definitive link between the metabolic stability of AldB (via its modification by LFTR) and YiaY or a specific cellular stress has yet to be elucidated.

Given the identification of T_2_-AldB as a substrate of LFTR, we considered the possibility that (under different conditions), MetAP cleavage of other proteins with the Nt sequence MTN, may generate additional LFTR substrates. Therefore, we searched the *E. coli* genome for sequences encoding proteins with the Nt sequence, MTN. From this analysis we identified 20 proteins (9 of unknown function), in which the Nt sequence (MTN) is located within the cytosol (i.e. cytosolic proteins, single or multi-pass inner membrane proteins). Of these 20 proteins, 12 lacked acidic residue near the N-terminus and hence were selected as potential LFTR-substrates (Supplementary Table 5). Interestingly, six of the proteins are known to be cleaved (at least partially) by MetAP and one protein (BarA) is a known ClpA-interacting protein (Butland et al., 2005;Rajagopala et al., 2014;Bienvenut et al., 2015) and hence may represent an additional Leu/N-degron substrate in *E. coli*. However, there is currently no direct evidence that any of these proteins are modified by LFTR. Therefore, more work is required to see if the metabolic stability of these proteins (including BarA) is influenced either by ClpS or LFTR.

To further examine the acceptor specificity of LFTR we used peptide arrays. This analysis demonstrated that residue(s) downstream of the N_d2_ can influence the specificity of LFTR. Based on our conjugation data, we propose a relative classification (favored, accepted and disfavored) for residues in position 2 of the acceptor. Although the vast majority of residues (e.g. small polar and small hydrophobic residues) are accepted in position 2 of the substrate (> 50% conjugation, relative to R-b, where b = R or K), only basic residues (i.e. Arg and Lys) are strongly favored in this position. In contrast, acidic residues are strongly disfavored in position 2 of the acceptor (< 20 % conjugation, relative to R-b,), while aromatic residues and Cys are also disfavored (< 30% conjugation, relative to R-b). Significantly, these direct conjugation data are not only generally consistent with the physicochemical properties of the LFTR binding pocket (Suto et al., 2006;Watanabe et al., 2007), but they are also highly consistent with the findings of Soffer, who examined the ability of select b-X dipeptides (where b = Arg or Lys and X = selected amino acids) to inhibit the LFTR-dependent conjugation of [^14^C]-Phe to α_S1_-casein (Soffer, 1973). Fittingly, the acceptor specificity of LFTR is comparable to the substrate specificity of the two other main components of the bacterial Leu/N-degron pathway, MetAP (Frottin et al., 2006) and the N-recognin, ClpS (Erbse et al., 2006;Wang et al., 2008b;Schuenemann et al., 2009), both of which disfavor acidic residues near the N-terminus of their substrates.

## AUTHOR CONTRIBUTIONS

Conceptualization, D.A.D. and K.N.T.; methodology, K.N.T. and D.A.D.; investigation, R.D.O and R.L.N.; writing original draft, R.D.O., D.A.D. and K.N.T.; supervision, project administration and funding acquisition, D.A.D. and K.N.T.

## ACKNOWLEDGEMENTS

We thank the Australian Proteome Analysis Facility (APAF) for performing Edman degradation, the National BioResource Project (NIG, Japan) for the *E. coli* strains used in this study.

## REFERENCES

Alhuwaider, A.H., and Dougan, D.A. (2017). AAA+ Machines of Protein Destruction in Mycobacteria. Front Mol Biosci 4, 49.

Bienvenut, W.V., Giglione, C., and Meinnel, T. (2015). Proteome-wide analysis of the amino terminal status of Escherichia coli proteins at the steady-state and upon deformylation inhibition. Proteomics 15, 2503–2518.

Bouchnak, I., and Van Wijk, K.J. (2019). N-Degron Pathways in Plastids. Trends Plant Sci 24, 917–926.

Butland, G., Peregrin-Alvarez, J.M., Li, J., Yang, W., Yang, X., Canadien, V., et al. (2005). Interaction network containing conserved and essential protein complexes in Escherichia coli. Nature 433, 531–537.

Catanzariti, A.M., Soboleva, T.A., Jans, D.A., Board, P.G., and Baker, R.T. (2004). An efficient system for high-level expression and easy purification of authentic recombinant proteins. Protein Sci 13, 1331–1339.

Chen, S.J., Wu, X., Wadas, B., Oh, J.H., and Varshavsky, A. (2017). An N-end rule pathway that recognizes proline and destroys gluconeogenic enzymes. Science 355.

Dissmeyer, N., Rivas, S., and Graciet, E. (2018). Life and death of proteins after protease cleavage: protein degradation by the N-end rule pathway. New Phytol 218, 929–935.

Dougan, D.A., Micevski, D., and Truscott, K.N. (2012). The N-end rule pathway: from recognition by N-recognins, to destruction by AAA+proteases. Biochim Biophys Acta 1823, 83–91.

Dougan, D.A., Mogk, A., Zeth, K., Turgay, K., and Bukau, B. (2002a). AAA+ proteins and substrate recognition, it all depends on their partner in crime. FEBS Lett 529, 6–10.

Dougan, D.A., Reid, B.G., Horwich, A.L., and Bukau, B. (2002b). ClpS, a substrate modulator of the ClpAP machine. Mol Cell 9, 673–683.

Dougan, D.A., Truscott, K.N., and Zeth, K. (2010). The bacterial N-end rule pathway: expect the unexpected. Mol Microbiol 76, 545–558.

Dougan, D.A., and Varshavsky, A. (2018). Understanding the Pro/N-end rule pathway. Nat Chem Biol 14, 415–416.

Erbse, A., Schmidt, R., Bornemann, T., Schneider-Mergener, J., Mogk, A., Zahn, R., et al. (2006). ClpS is an essential component of the N-end rule pathway in Escherichia coli. Nature 439, 753–756.

Flynn, J.M., Neher, S.B., Kim, Y.I., Sauer, R.T., and Baker, T.A. (2003). Proteomic discovery of cellular substrates of the ClpXP protease reveals five classes of ClpX-recognition signals. Mol Cell 11, 671–683.

Franchini, A.G., and Egli, T. (2006). Global gene expression in Escherichia coli K-12 during short-term and long-term adaptation to glucose-limited continuous culture conditions. Microbiology (Reading) 152, 2111–2127.

Frottin, F., Martinez, A., Peynot, P., Mitra, S., Holz, R.C., Giglione, C., et al. (2006). The proteomics of N-terminal methionine cleavage. Mol Cell Proteomics 5, 2336–2349.

Gao, X., Yeom, J., and Groisman, E.A. (2019). The expanded specificity and physiological role of a widespread N-degron recognin. Proc Natl Acad Sci U S A 116, 18629–18637.

Geissler, A., Chacinska, A., Truscott, K.N., Wiedemann, N., Brandner, K., Sickmann, A., et al. (2002). The mitochondrial presequence translocase: an essential role of Tim50 in directing preproteins to the import channel. Cell 111, 507–518.

Gibbs, D.J., Bacardit, J., Bachmair, A., and Holdsworth, M.J. (2014). The eukaryotic N-end rule pathway: conserved mechanisms and diverse functions. Trends Cell Biol 24, 603–611.

Graciet, E., Hu, R.G., Piatkov, K., Rhee, J.H., Schwarz, E.M., and Varshavsky, A. (2006). Aminoacyl-transferases and the N-end rule pathway of prokaryotic/eukaryotic specificity in a human pathogen. Proc Natl Acad Sci U S A 103, 3078–3083.

Gur, E., Ottofueling, R., and Dougan, D.A. (2013). Machines of destruction - AAA+ proteases and the adaptors that control them. Subcell Biochem 66, 3–33.

Ho, K.K., and Weiner, H. (2005). Isolation and characterization of an aldehyde dehydrogenase encoded by the aldB gene of Escherichia coli. J Bacteriol 187, 1067–1073.

Humbard, M.A., Surkov, S., De Donatis, G.M., Jenkins, L.M., and Maurizi, M.R. (2013). The N-degradome of Escherichia coli: limited proteolysis in vivo generates a large pool of proteins bearing N-degrons. J Biol Chem 288, 28913–28924.

Hwang, C.S., Shemorry, A., and Varshavsky, A. (2010). N-terminal acetylation of cellular proteins creates specific degradation signals. Science 327, 973–977.

Kaji, A., Kaji, H., and Novelli, G.D. (1963). A soluble amino acid incorporating system. Biochem Biophys Res Commun 10, 406–409.

Kawai, Y., Matsumoto, S., Ling, Y., Okuda, S., and Tsuneda, S. (2018). AldB controls persister formation in Escherichia coli depending on environmental stress. Microbiol Immunol 62, 299–309.

Keiler, K.C., Waller, P.R., and Sauer, R.T. (1996). Role of a peptide tagging system in degradation of proteins synthesized from damaged messenger RNA. Science 271, 990–993.

Kim, H.K., Kim, R.R., Oh, J.H., Cho, H., Varshavsky, A., and Hwang, C.S. (2014). The N-terminal methionine of cellular proteins as a degradation signal. Cell 156, 158–169.

Kirstein, J., Moliere, N., Dougan, D.A., and Turgay, K. (2009). Adapting the machine: adaptor proteins for Hsp100/Clp and AAA+ proteases. Nat Rev Microbiol 7, 589–599.

Kress, W., Mutschler, H., and Weber-Ban, E. (2009). Both ATPase domains of ClpA are critical for processing of stable protein structures. J Biol Chem 284, 31441–31452.

Leibowitz, M.J., and Soffer, R.L. (1970). Enzymatic modification of proteins. 3. Purification and properties of a leucyl, phenylalanyl transfer ribonucleic acid protein transferase from Escherichia coli. J Biol Chem 245, 2066–2073.

Lucas, X., and Ciulli, A. (2017). Recognition of substrate degrons by E3 ubiquitin ligases and modulation by small-molecule mimicry strategies. Curr Opin Struct Biol 44, 101–110.

Ma, F., Wang, T., Ma, X., and Wang, P. (2014). Identification and characterization of protein encoded by orf382 as L-threonine dehydrogenase. J Microbiol Biotechnol 24, 748–755.

Mahmoud, S.A., and Chien, P. (2018). Regulated Proteolysis in Bacteria. Annu Rev Biochem 87, 677–696.

Mogk, A., Schmidt, R., and Bukau, B. (2007). The N-end rule pathway for regulated proteolysis: prokaryotic and eukaryotic strategies. Trends Cell Biol 17, 165–172.

Neuwald, A.F., Aravind, L., Spouge, J.L., and Koonin, E.V. (1999). AAA+: A class of chaperone-like ATPases associated with the assembly, operation, and disassembly of protein complexes. Genome Res 9, 27–43.

Newman, E.B., Kapoor, V., and Potter, R. (1976). Role of L-threonine dehydrogenase in the catabolism of threonine and synthesis of glycine by Escherichia coli. J Bacteriol 126, 1245–1249.

Nguyen, K.T., Kim, J.M., Park, S.E., and Hwang, C.S. (2019). N-terminal methionine excision of proteins creates tertiary destabilizing N-degrons of the Arg/N-end rule pathway. J Biol Chem 294, 4464–4476.

Nguyen, K.T., Mun, S.H., Lee, C.S., and Hwang, C.S. (2018). Control of protein degradation by N-terminal acetylation and the N-end rule pathway. Exp Mol Med 50, 1–8.

Ninnis, R.L., Spall, S.K., Talbo, G.H., Truscott, K.N., and Dougan, D.A. (2009). Modification of PATase by L/F-transferase generates a ClpS-dependent N-end rule substrate in Escherichia coli. EMBO J 28, 1732–1744.

Ogura, T., and Wilkinson, A.J. (2001). AAA+ superfamily ATPases: common structure--diverse function. Genes Cells 6, 575–597.

Piatkov, K.I., Vu, T.T., Hwang, C.S., and Varshavsky, A. (2015). Formyl-methionine as a degradation signal at the N-termini of bacterial proteins. Microb Cell 2, 376–393.

Rajagopala, S.V., Sikorski, P., Kumar, A., Mosca, R., Vlasblom, J., Arnold, R., et al. (2014). The binary protein-protein interaction landscape of Escherichia coli. Nat Biotechnol 32, 285–290.

Rivera-Rivera, I., Roman-Hernandez, G., Sauer, R.T., and Baker, T.A. (2014). Remodeling of a delivery complex allows ClpS-mediated degradation of N-degron substrates. Proc Natl Acad Sci U S A 111, E3853–3859.

Roman-Hernandez, G., Hou, J.Y., Grant, R.A., Sauer, R.T., and Baker, T.A. (2011). The ClpS adaptor mediates staged delivery of N-end rule substrates to the AAA+ ClpAP protease. Mol Cell 43, 217–228.

Sauer, R.T., and Baker, T.A. (2011). AAA+ proteases: ATP-fueled machines of protein destruction. Annu Rev Biochem 80, 587–612.

Scarpulla, R.C., Deutch, C.E., and Soffer, R.L. (1976). Transfer of methionyl residues by leucyl, phenylalanyl-tRNA-protein transferase. Biochem Biophys Res Commun 71, 584–589.

Schmidt, R., Zahn, R., Bukau, B., and Mogk, A. (2009). ClpS is the recognition component for Escherichia coli substrates of the N-end rule degradation pathway. Mol Microbiol 72, 506–517.

Schuenemann, V.J., Kralik, S.M., Albrecht, R., Spall, S.K., Truscott, K.N., Dougan, D.A., et al. (2009). Structural basis of N-end rule substrate recognition in Escherichia coli by the ClpAP adaptor protein ClpS. EMBO Rep 10, 508–514.

Shemorry, A., Hwang, C.S., and Varshavsky, A. (2013). Control of protein quality and stoichiometries by N-terminal acetylation and the N-end rule pathway. Mol Cell 50, 540–551.

Shrader, T.E., Tobias, J.W., and Varshavsky, A. (1993). The N-end rule in Escherichia coli: cloning and analysis of the leucyl, phenylalanyl-tRNA-protein transferase gene aat. J Bacteriol 175, 4364–4374.

Soffer, R.L. (1973). Peptide acceptors in the arginine transfer reaction. J Biol Chem 248, 2918–2921.

Striebel, F., Kress, W., and Weber-Ban, E. (2009). Controlled destruction: AAA+ ATPases in protein degradation from bacteria to eukaryotes. Curr Opin Struct Biol 19, 209–217.

Suto, K., Shimizu, Y., Watanabe, K., Ueda, T., Fukai, S., Nureki, O., et al. (2006). Crystal structures of leucyl/phenylalanyl-tRNA-protein transferase and its complex with an aminoacyl-tRNA analog. EMBO J 25, 5942–5950.

Tasaki, T., Sriram, S.M., Park, K.S., and Kwon, Y.T. (2012). The N-end rule pathway. Annu Rev Biochem 81, 261–289.

Timms, R.T., and Koren, I. (2020). Tying up loose ends: the N-degron and C-degron pathways of protein degradation. Biochem Soc Trans 48, 1557–1567.

Tobias, J.W., Shrader, T.E., Rocap, G., and Varshavsky, A. (1991). The N-end rule in bacteria. Science 254, 1374–1377.

Tu, G.F., Reid, G.E., Zhang, J.G., Moritz, R.L., and Simpson, R.J. (1995). C-terminal extension of truncated recombinant proteins in Escherichia coli with a 10Sa RNA decapeptide. J Biol Chem 270, 9322–9326.

Varshavsky, A. (2011). The N-end rule pathway and regulation by proteolysis. Protein Sci 20, 1298–1345.

Varshavsky, A. (2019). N-degron and C-degron pathways of protein degradation. Proc Natl Acad Sci U S A 116, 358–366.

Wang, K.H., Oakes, E.S., Sauer, R.T., and Baker, T.A. (2008a). Tuning the strength of a bacterial N-end rule degradation signal. J Biol Chem 283, 24600–24607.

Wang, K.H., Roman-Hernandez, G., Grant, R.A., Sauer, R.T., and Baker, T.A. (2008b). The molecular basis of N-end rule recognition. Mol Cell 32, 406–414.

Watanabe, K., Toh, Y., Suto, K., Shimizu, Y., Oka, N., Wada, T., et al. (2007). Protein-based peptide-bond formation by aminoacyl-tRNA protein transferase. Nature 449, 867–871.

Xu, J., and Johnson, R.C. (1995). aldB, an RpoS-dependent gene in Escherichia coli encoding an aldehyde dehydrogenase that is repressed by Fis and activated by Crp. J Bacteriol 177, 3166–3175.

Yang, C.I., Hsieh, H.H., and Shan, S.O. (2019). Timing and specificity of cotranslational nascent protein modification in bacteria. Proc Natl Acad Sci U S A 116, 23050–23060.

Yeom, J., and Groisman, E.A. (2019). Activator of one protease transforms into inhibitor of another in response to nutritional signals. Genes Dev 33, 1280–1292.

Yeom, J., Wayne, K.J., and Groisman, E.A. (2017). Sequestration from Protease Adaptor Confers Differential Stability to Protease Substrate. Mol Cell 66, 234–246 e235.

