## Supplementary material for "Novel modification by L/F-tRNA-protein transferase (LFTR) generates a Leu/N-degron ligand in *Escherichia coli*": S

### Supplementary Figures and Tables

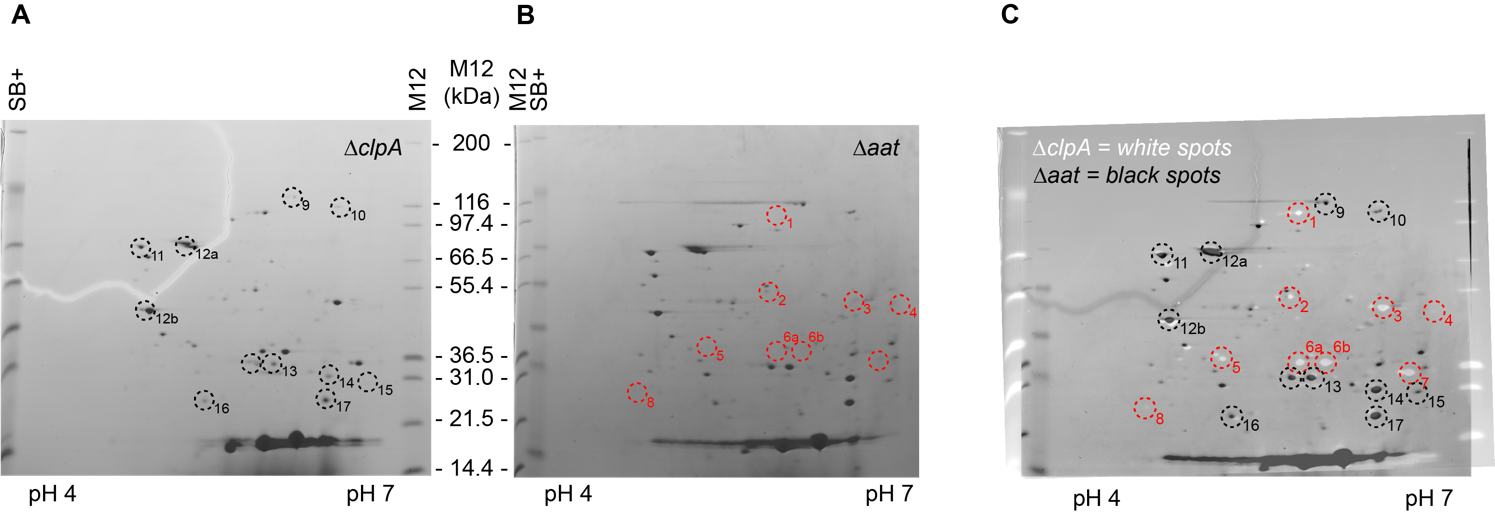

**Supplementary Figure 1.** *E. coli* N-degron bearing proteins from (**A**) ∆*clpA* and (**B**) ∆*aat* cells were specifically eluted from immobilized ClpS using FR dipeptide and analyzed by 2D-PAGE. (**C**) To visualize the difference between the two elution profiles, the image for ∆*clpA* was inverted (proteins = white spots) and overlayed with ∆*aat* cells. Proteins not recovered from ∆*aat* cells (i.e. LFTR-dependent N-degrons) are numbered 1 – 8 (red) and those recovered from both ∆*clpA* and ∆*aat* are numbered 9 – 17 (black).

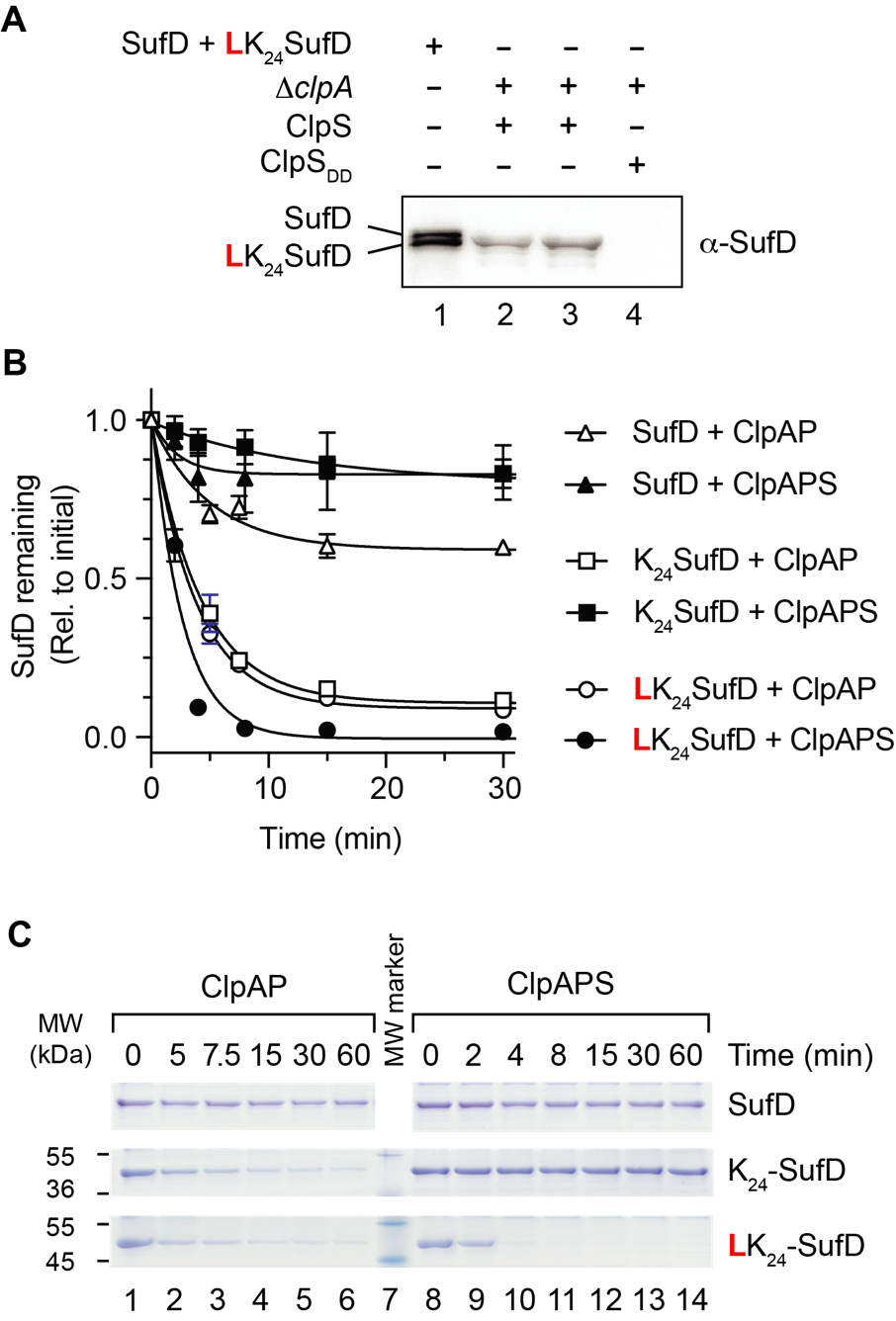

**Supplementary Figure 2.**

(**A**) N-degrons ligands from ∆*clpA* *E. coli* cells were specifically eluted (using FR dipeptide) from either immobilized wild type ClpS (lanes 2 and 3) or mutant ClpS_DD_ (lane 4) and compared to recombinant SufD (lane 1, upper band) and LK_24_SufD (lane 1, lower band) using a specific α-SufD antisera. (**B**) The turnover of recombinant SufD (triangles), K_24_SufD (squares) and LK_24_SufD (circles) was monitored in the presence of ClpAP (open symbols) or ClpAPS (filled symbols). Error bars represent the standard error of the mean (S.E.M.) from the quantitation of 3 independent experiments (**C**) 16.5% Tris-Tricine SDS-PAGE showing a representative assay for the turnover of recombinant protein substrates by ClpAP (lanes 1 – 6) or ClpAPS (lanes 8 – 14), MW marker is shown in lane 7.

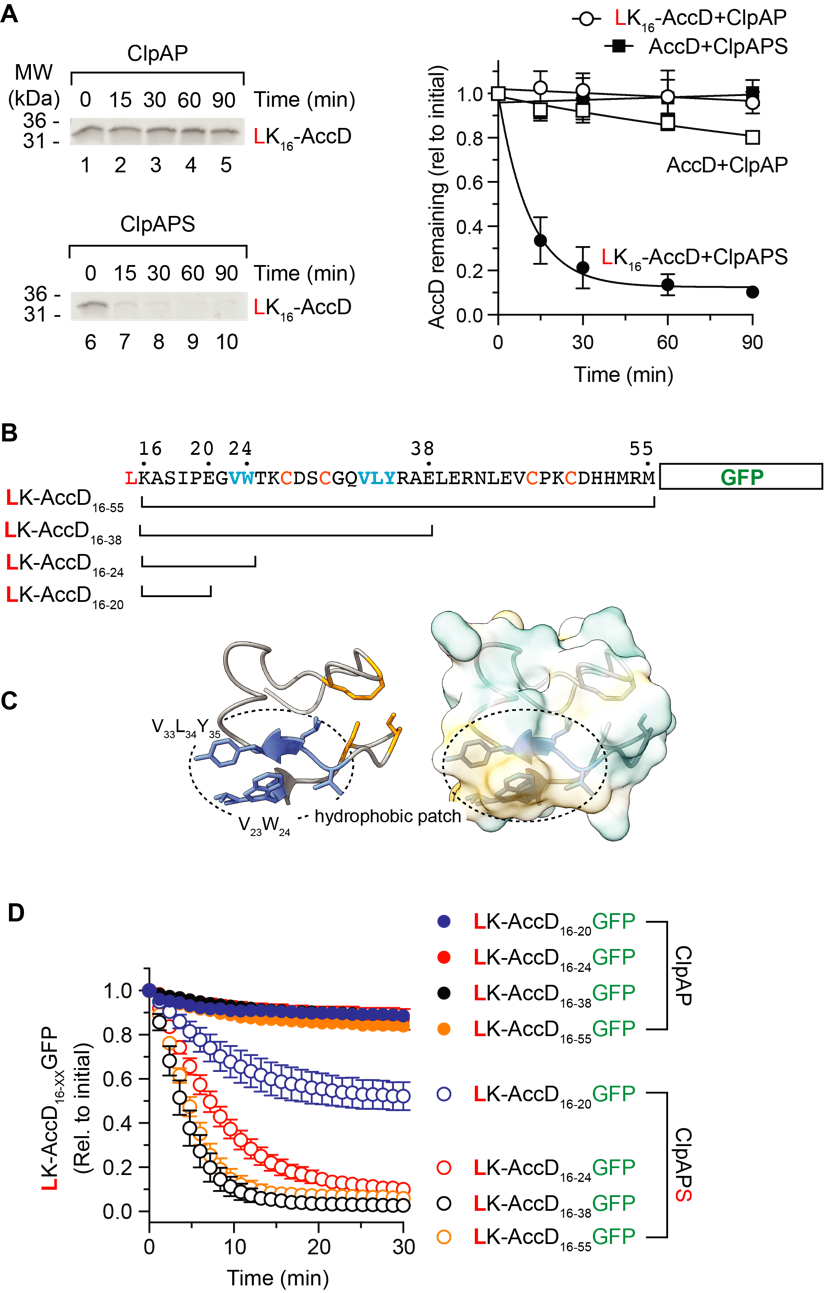

**Supplementary Figure 3.**

(**A,** Right panel) The turnover of recombinant AccD (squares) and LK_16_AccD (circles) was monitored in the presence of ClpAP (open symbols) or ClpAPS (filled symbols). Error bars represent the standard error of the mean (S.E.M.) from the quantitation of 3 independent experiments (**A,** Left panel) 16.5% Tris-Tricine SDS-PAGE showing a representative assay for the turnover of recombinant LK_16_AccD by ClpAP (lanes 1 – 5) or ClpAPS (lanes 6 – 10), MW marker are indicated. (**B**) Schematic representation of the GFP fusions used in (D). (**C**) Structural mode of AccD N-terminal region (PDB: 2F9Y) (residues 23 – 55) shown in ribbon (left panel) and surface representation (right panel), highlighting the hydrophobic residues (blue) and conserved Cys residues (yellow). (**D**) The ClpAP-dependent turnover of LK-AccD_16-20_GFP (blue symbols), LK-AccD_16-24_GFP (red symbols), LK-AccD_16-38_GFP (red symbols) and LK-AccD_16-55_GFP (orange symbols) was monitored by a change in fluorescence in the absence (filled symbols) or presence (open symbols) of ClpS.

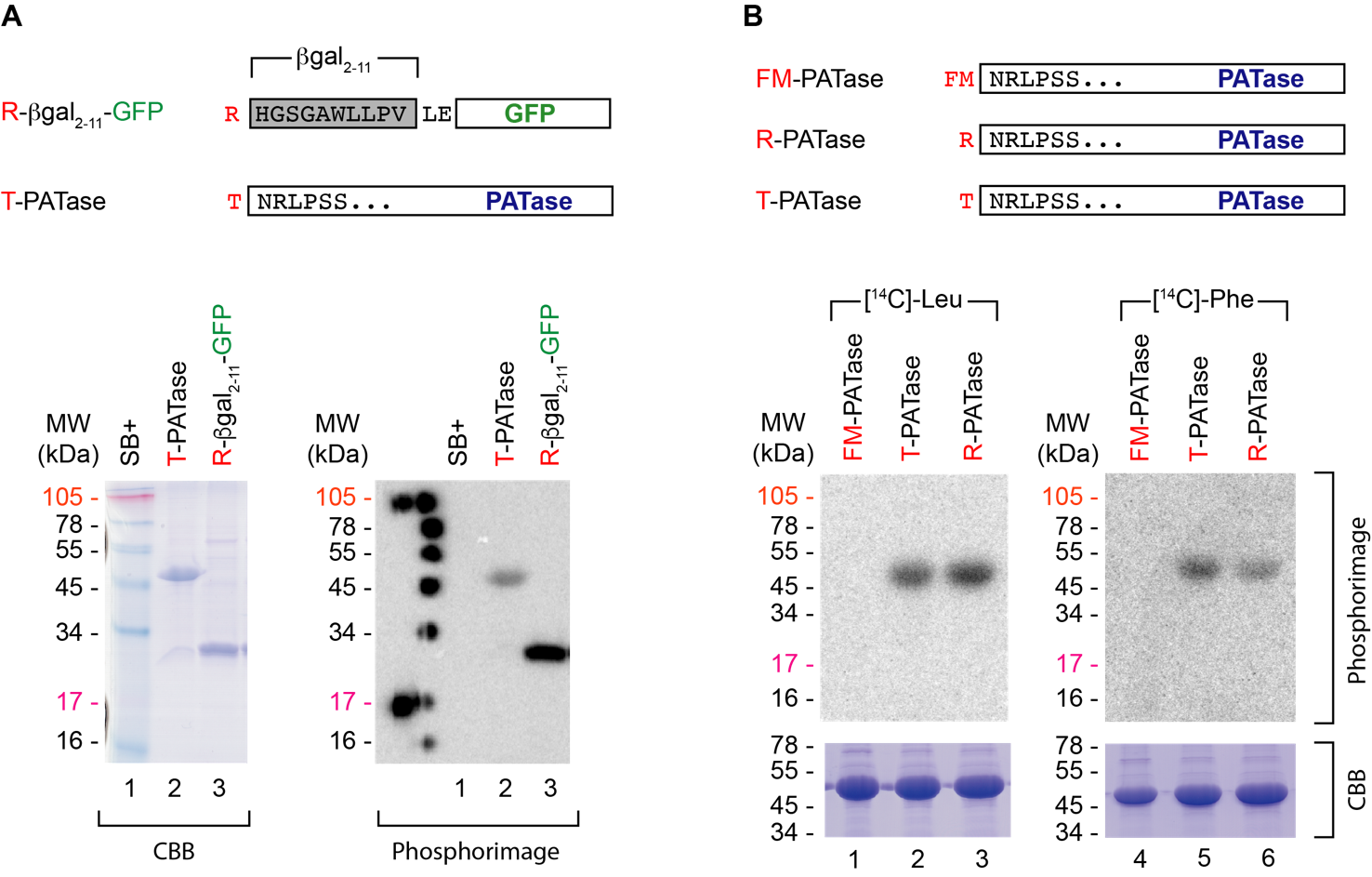

**Supplementary Figure 4.**

The Nt conjugation of T-PATase and R-β-gal_2-10_-GFP *in vitro* is dependent on LFTR. (**A**, Upper panel) Schematic representation of R-β-gal_2-10_-GFP and T-PATase. (**A**, Lower panel) Recombinant T-PATase and R-β-gal_2-10_-GFP were separated by 12.5% SDS-PAGE and then stained with Coomassie Brilliant Blue (CBB, left panel). Following drying of the polyacrylamide gel, the [^14^C]-Leu radiolabelled proteins were detected by phosphor image analysis using a Typhoon Trio Variable Mode Imager (right panel). (**B,** Upper panel) Schematic representation of various recombinant PATase fusion proteins (FM-PATase, R-PATase and T-PATase). X-PATase was separated by 12.5% SDS-PAGE and then stained with Coomassie Brilliant Blue (CBB, lower gel image). Following drying of the polyacrylamide gel, the [^14^C]-Leu (left) or [^14^C]-Phe radiolabelled proteins were detected by phosphor image analysis using a Typhoon Trio Variable Mode Imager (Phosphorimage, upper gel image).

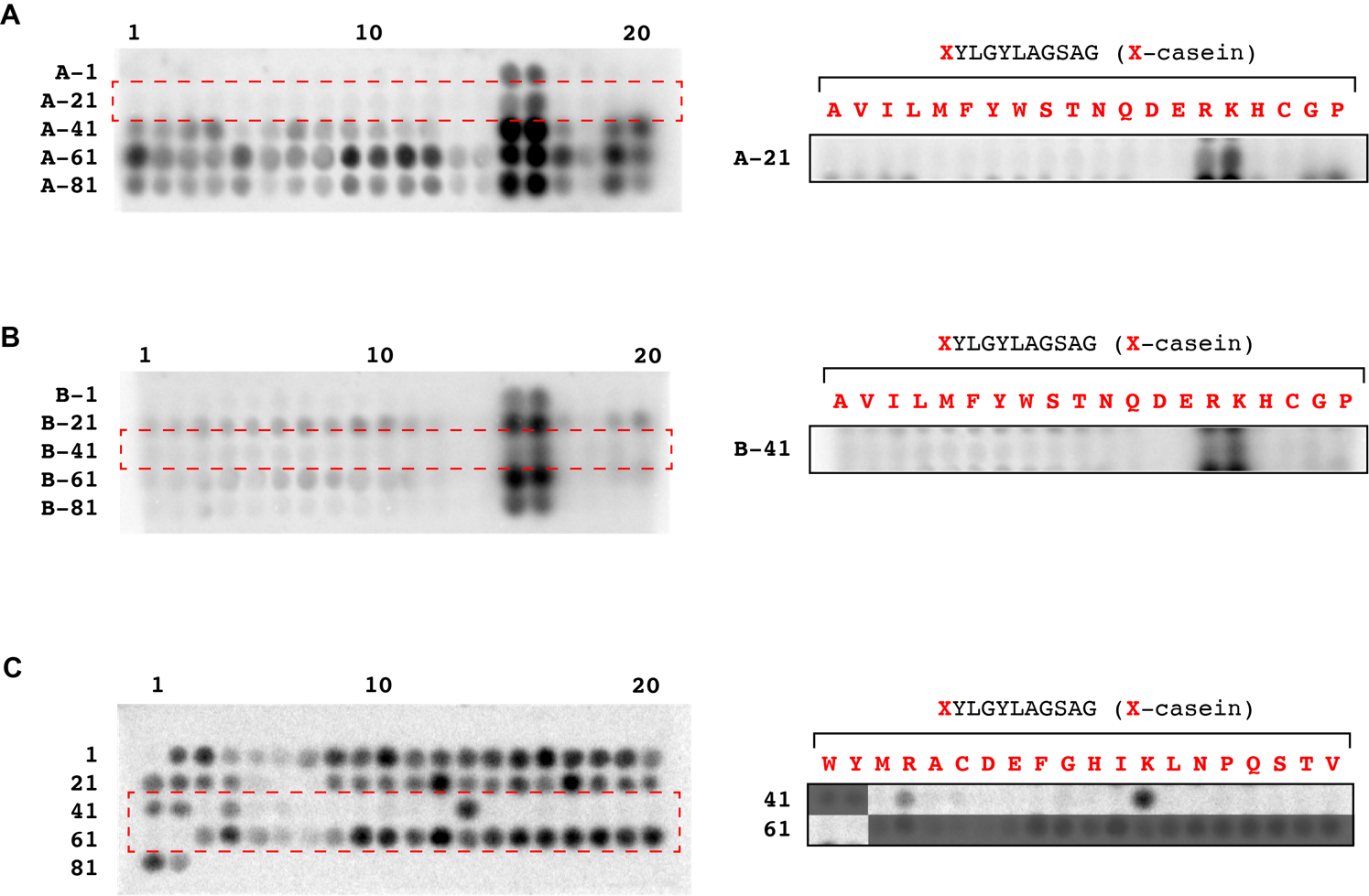

**Supplementary Figure 5.**

[^14^C]-Leu phosphorimages of 11-mer peptide libraries (**A**), (**B**) and (**C**) with LFTR. X-casein peptide sport are indicated by a red dotted box. Peptide sequences for X-casein peptides are indicated (right panel). For details of all other peptide sequences see Supplementary Tables 2 - 4.

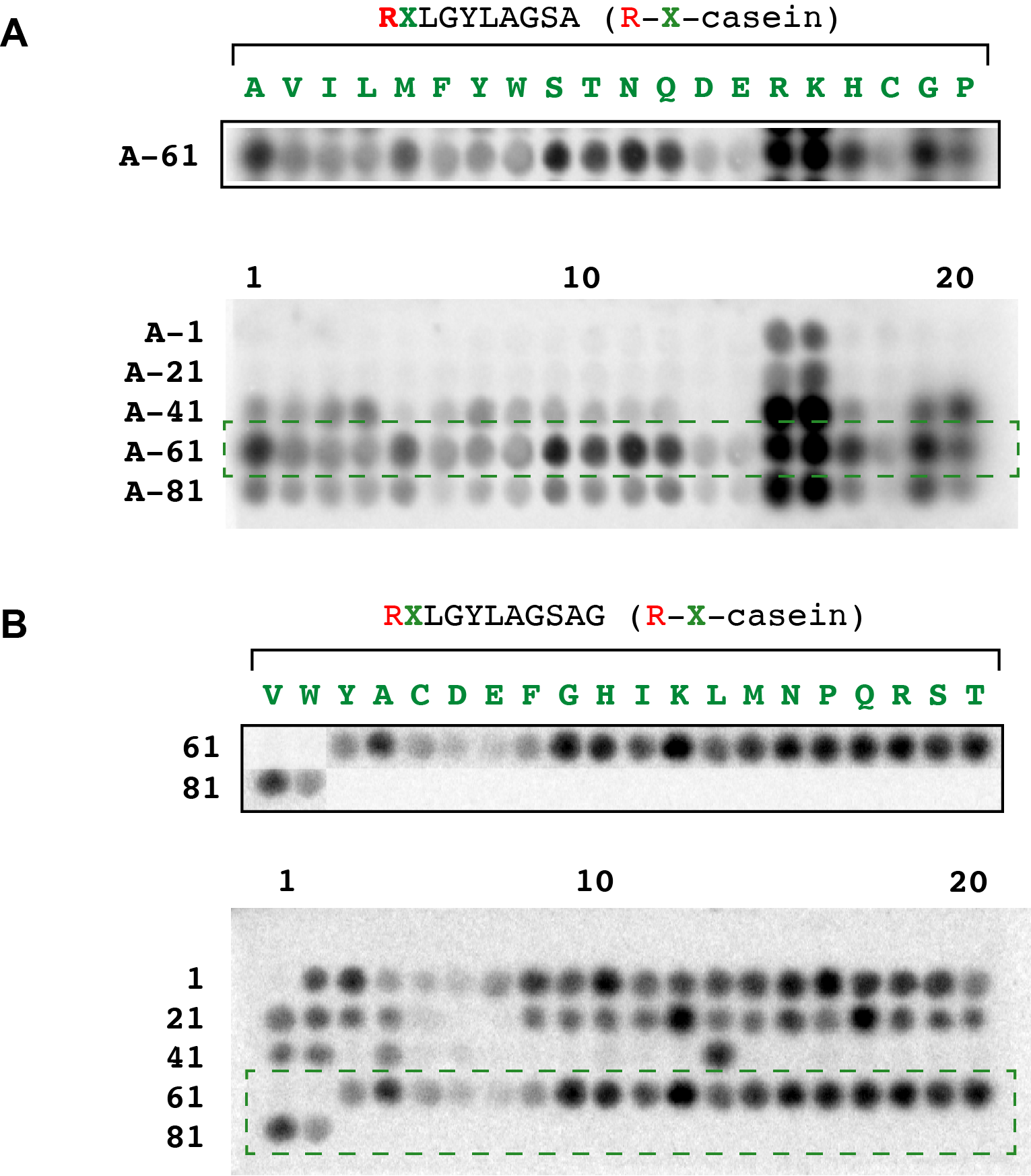

**Supplementary Figure 6.**

[^14^C]-Leu phosphorimages of 11-mer peptide libraries (**A**) and (**B**) with LFTR highlighting R-X-casein peptide spots (green dotted box). Peptide sequences for R-X-casein peptides are indicated below each peptide library panel. For details of all other peptide sequences see Supplementary Tables 2 and 3.

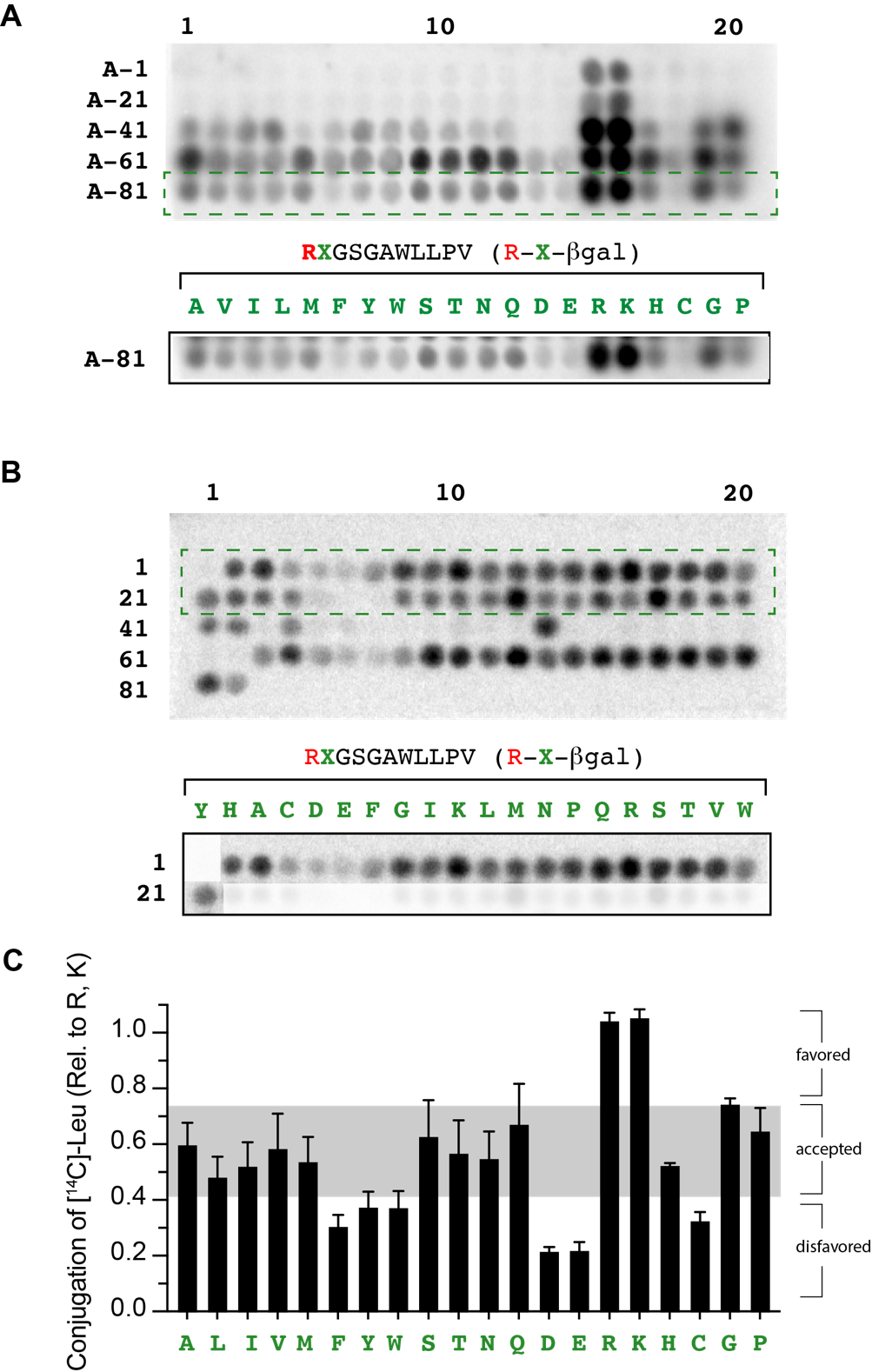

**Supplementary Figure 7.**

[^14^C]-Leu phosphorimages of 11-mer peptide libraries (**A**) and (**B**) with LFTR highlighting R-X-βgal peptide spots (green dotted box). Peptide sequences for R-X-βgal peptides are indicated below each peptide library panel. For details of all other peptide sequences see Supplementary Tables 2 and 3. (**C**) Conjugation of [^14^C]-Leu to R-X-βgal, relative to the average conjugation of R-b-βgal, where b = R, K. Relative conjugation activity was determined from two independent experiments and is separated into three broad categories (favored (>75%), accepted (40 – 75%, grey panel) and disfavored (< 40%)).

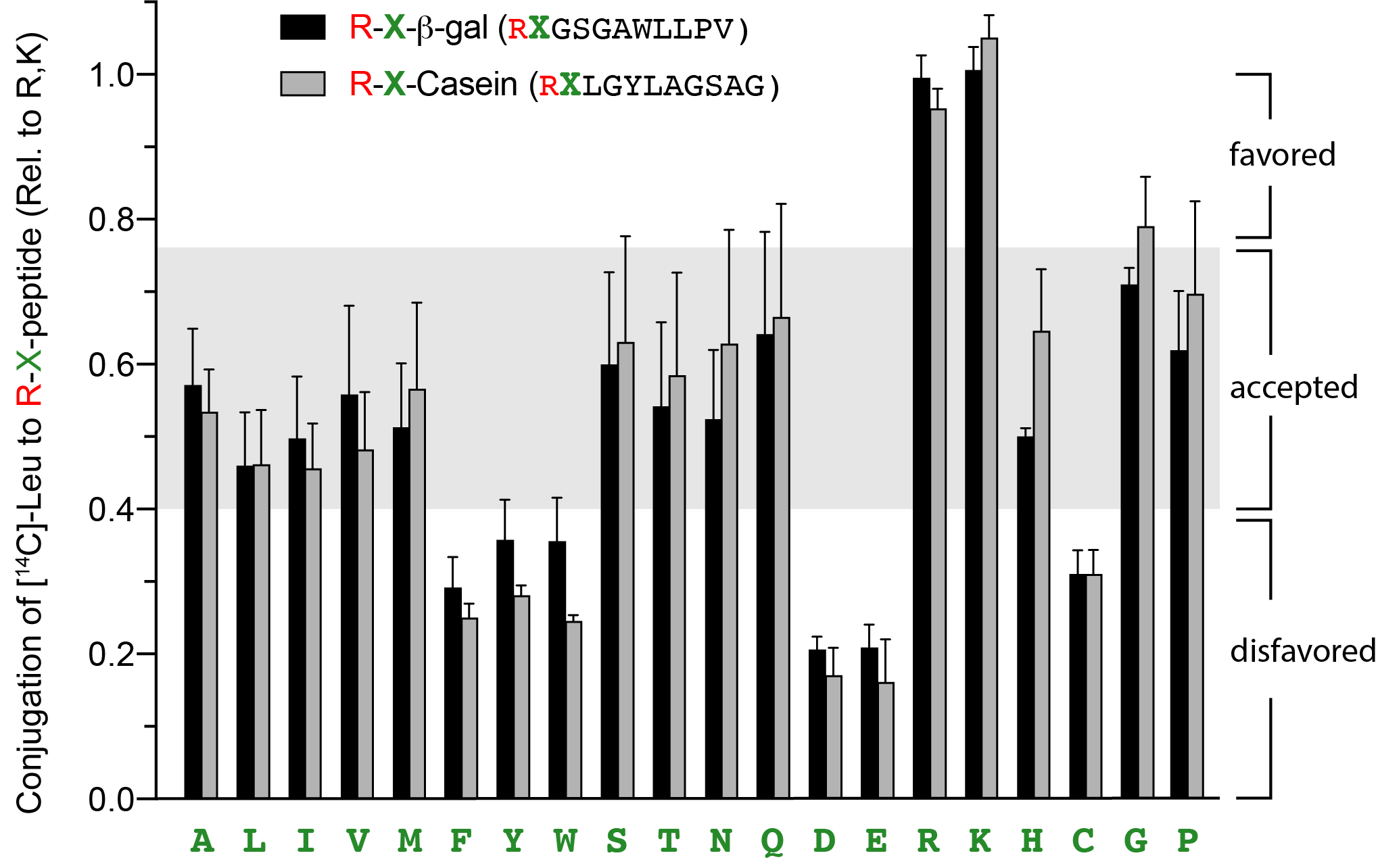

**Supplementary Figure 8.**

Comparison of the relative LFTR-dependent conjugation activity of R-X-βgal (black bars) and R-X-casein (grey bars) as illustrated in Supplementary Figures 6 and 7. Relative conjugation activity is separated into three broad categories (favored (>75%), accepted (40 – 75%, grey panel) and disfavored (< 40%)).

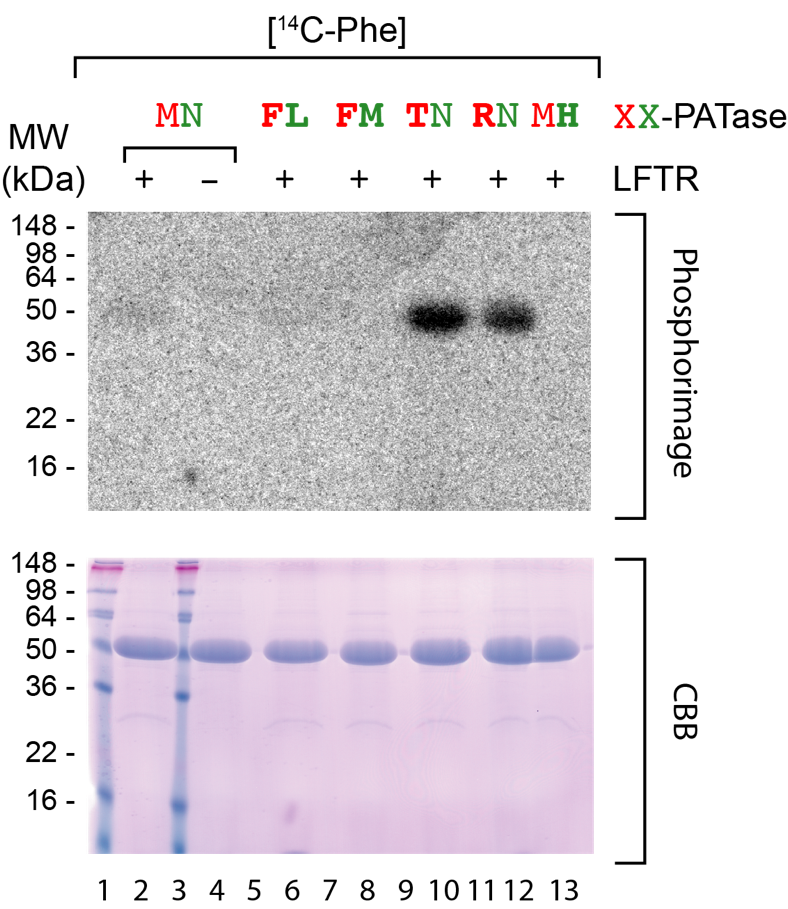

**Supplementary Figure 9.**

The Nt conjugation of XX-PATase *in vitro* is dependent on LFTR. Recombinant XX-PATase were separated by 12.5% SDS-PAGE and then stained with Coomassie Brilliant Blue (CBB, lower panel). As a control MN-PATase was incubated in the presence (lane 2) and absence (lane 4) of LFTR. All other XX-PATase variants were incubated in the presence of LFTR. FL-PATase (lane 6), FM-PATase (lane 8) TN-PATase (lane 10) RN-PATase (lane 12) and MH-PATase (lane 13). See blue + MW markers (Lanes 1 and 3). Lanes 5, 7, 9 and 11 lacked proteins. Following drying of the polyacrylamide gel, the [^14^C]-Phe radiolabeled proteins were detected by phosphor image analysis using a Typhoon Trio Variable Mode Imager (upper panel).

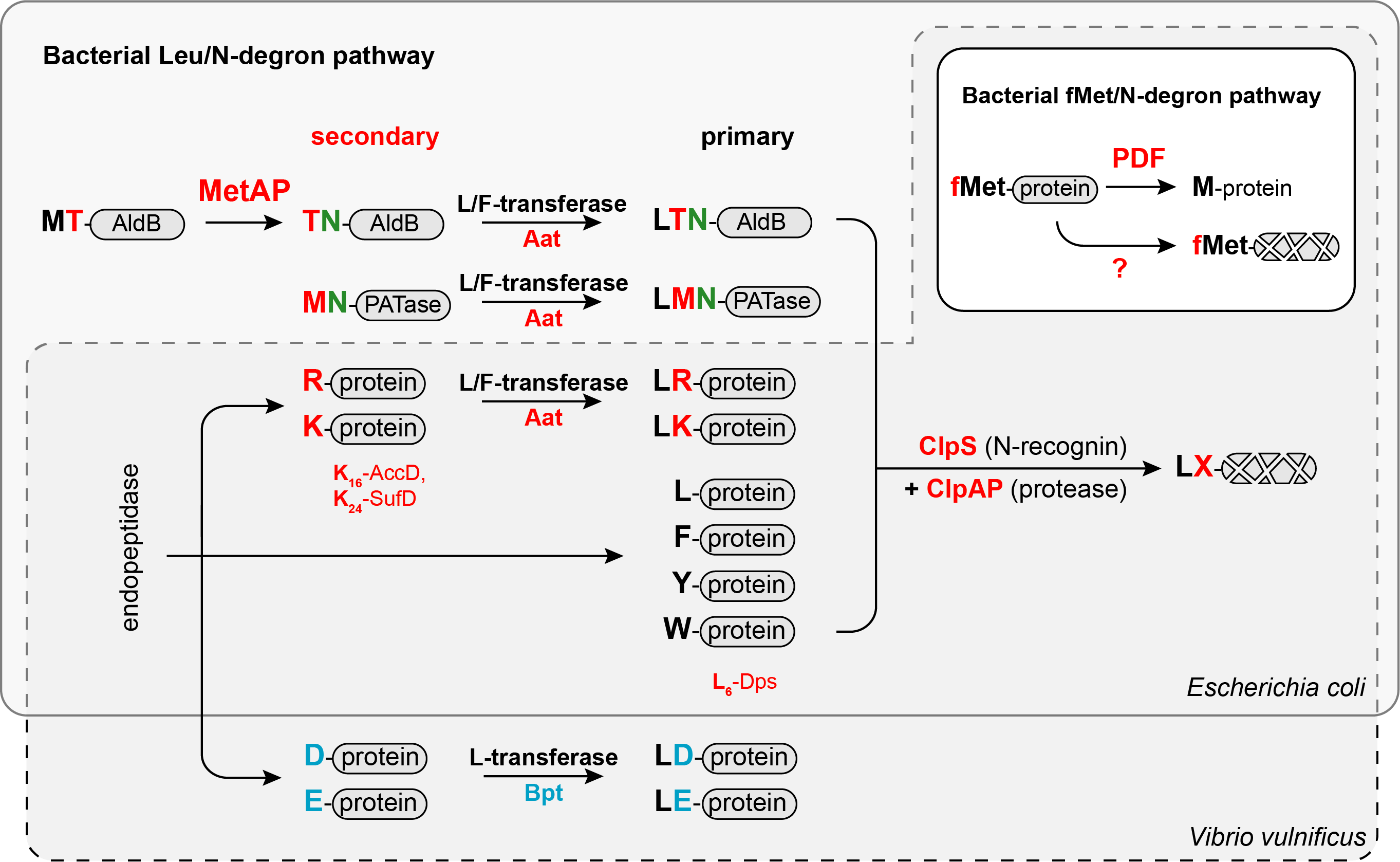

**Supplementary Figure 10.**

**The bacterial N-end rule pathways.** The fMet/N-degron pathway is responsible for the quality control of proteins that retain the formyl group as a result of incorrect processing by peptide deformylase (PDF). The N-recognin and/or protease responsible for the recognition and degradation of fMet/N-degrons is currently unknown. The Leu/N-degron pathway is the principal N-degron pathway in bacteria. N-terminal (Nt) destabilizing (Nd) residues are either primary or secondary. Primary Nd residues (L, F, Y and W) are recognized by the N-recognin, ClpS and delivered to the ATP-dependent protease, ClpAP. In *Vibrio vulnificus* primary Nd residues are transferred to secondary Nd residues by two different enzymes. Primary Nd residues (L or F) are transferred by L/F-transferase (LFTR) to basic secondary Nd residues (R and K), while the Nd residue (L) is transferred to acidic secondary Nd residues (D and E), by the L-transferase Bpt (Bacterial protein transferase). In *E. coli* primary Nd residues (L or F) are transferred by L/F-transferase to basic secondary Nd residues (R and K). In addition to basic secondary Nd residues, the initiating Met (of PATase) is recognized by L/F-transferase. While in the case of AldB, the Nt Thr (following excision of the initiating Met, by MetAP) is recognized as a secondary destabilizing residue by L/F-transferase. Endoproteolytic processing (by unidentified proteases) generates protein fragments bearing either a primary (e.g. L_6_-Dps) or secondary (e.g. K_16_-AccD, K_24_-SufD) destabilizing residue.

**SUPPLEMENTARY TABLE 1 – Putative L/F-transferase substrates**

| **Protein (Spot Ref.)** | **Protein Name** | **Accession number** | **MW (kDa)** | **Function** | **N-terminal Sequence** |
| --- | --- | --- | --- | --- | --- |
| **1** | ClpB |  | ~100 | AAA+ chaperone | n.d. |
| **2** | AldB | P37685 | 57 | Catalyzes the NADP-dependent oxidation of diverse aldehydes. | L(F)-T_2_NNP |
| **3** | PATase | P42588 | 50 | Putrescine aminotransferase | L(F)-M_1_NRLP |
| **4** | SufD | P77689 | 47 | Part of the SufBCD complex – assembly or repair of Fe-S clusters under oxidative stress | L(F)-K_24_RSPQA^¥^ |
| **5** | RsgA (EngS) | P39286 | 36 | Small ribosomal subunit biogenesis GTPase RsgA | L-R_21_LKTSK^§^ |
| **6a, 6b** | AccA | P0ABD5 | 35 | alpha subunit of the Acetyl-CoA carboxylase (ACCase) complex. | S_2_LNFL |
| **7** | AccD | P0A9Q5 | 34 | beta subunit of the Acetyl-CoA carboxylase (ACCase) complex. | L-K_16_AS |
| **8** | SufC | P77499 | 27 | Part of the SufBCD complex – assembly or repair of Fe-S clusters under oxidative stress. | n.d. |
| **9** | PutA | P09546 | 140 | Bifunctional protein PutA | n.d. |
| **10** | Odo1 | P0AFG3 | 130 | 2-oxoglutarate dehydrogenase E1 component | n.d. |
| **11** | DnaK | P0A6Y8 | 70 | Hsp70 chaperone | n.d. |
| **12a, 12b** | KatG | P13029 | 80 | Catalase-peroxidase | n.d. |
| **13** | ? | - | ~33 | - | n.d. |
| **14** | UspE | P0AAC0 | 35 | Universal stress protein E | n.d. |
| **15** | YfiF | P0AGJ5 | 37 | Uncharacterized tRNA/rRNA methyltransferase | n.d. |
| **16** | CmoA | P76290 | 28 | Carboxy-S-adenosyl-L-methionine synthase | n.d. |
| **17** | AphA | P0AE22 | 24 | Class B acid phosphatase | L_24_ASSPS^†^ |
| **Dps** | Dps | P27430 | 19 | DNA protection during starvation protein. | L_6_VKSK |

^¥^ proposed by mobility on SDS-PAGE (Supplementary Figure 2)

^§^ (Daigle and Brown, 2004)

^†^ (Link et al., 1997)

**SUPPLEMENTARY TABLE 2** Spot sequences for peptide library **1**

```
Spot	sequence 	Spot	sequence
-----------------------	-----------------------
```

1. MHGSGAWLLPV	49.	FYLGYLAGSAG

```
2.	RHGSGAWLLPV	50.	GYLGYLAGSAG
3.	RAGSGAWLLPV	51.	HYLGYLAGSAG
4.	RCGSGAWLLPV	52.	IYLGYLAGSAG
5.	RDGSGAWLLPV	53.	KYLGYLAGSAG
6.	REGSGAWLLPV	54.	LYLGYLAGSAG
7.	RFGSGAWLLPV	55.	NYLGYLAGSAG
8.	RGGSGAWLLPV	56.	PYLGYLAGSAG
9.	RIGSGAWLLPV	57.	QYLGYLAGSAG
10.	RKGSGAWLLPV	58.	SYLGYLAGSAG
11.	RLGSGAWLLPV	59.	TYLGYLAGSAG
12.	RMGSGAWLLPV	60.	VYLGYLAGSAG
13.	RNGSGAWLLPV	61.	WYLGYLAGSAG
14.	RPGSGAWLLPV	62.	YYLGYLAGSAG
15.	RQGSGAWLLPV	63.	RYLGYLAGSAG
16.	RRGSGAWLLPV	64.	RALGYLAGSAG
17.	RSGSGAWLLPV	65.	RCLGYLAGSAG
18.	RTGSGAWLLPV	66.	RDLGYLAGSAG
19.	RVGSGAWLLPV	67	RELGYLAGSAG
20.	RWGSGAWLLPV	68.	RFLGYLAGSAG
21.	RYGSGAWLLPV	69.	RGLGYLAGSAG
22.	FNRLPSSASAL	70.	RHLGYLAGSAG
23.	MNRLPSSASAL	71.	RILGYLAGSAG
24.	MARLPSSASAL	72.	RKLGYLAGSAG
25.	MCRLPSSASAL	73.	RLLGYLAGSAG
26.	MDRLPSSASAL	74.	RMLGYLAGSAG
27.	MERLPSSASAL	75.	RNLGYLAGSAG
28.	MFRLPSSASAL	76.	RPLGYLAGSAG
29.	MGRLPSSASAL	77.	RQLGYLAGSAG
30.	MHRLPSSASAL	78.	RRLGYLAGSAG
31.	MIRLPSSASAL	79.	RSLGYLAGSAG
32.	MKRLPSSASAL	80.	RTLGYLAGSAG
33.	MLRLPSSASAL	81.	RVLGYLAGSAG
34.	MMRLPSSASAL	82.	RWLGYLAGSAG
35.	MPRLPSSASAL
36.	MQRLPSSASAL
37.	MRRLPSSASAL
38.	MSRLPSSASAL
39.	MTRLPSSASAL
40.	MVRLPSSASAL
41.	MWRLPSSASAL
42.	MYRLPSSASAL
43.	MYLGYLAGSAG
44.	RYLGYLAGSAG
45.	AYLGYLAGSAG
46.	CYLGYLAGSAG
47.	DYLGYLAGSAG
48.	EYLGYLAGSAG
```

**SUPPLEMENTARY TABLE 3** Spot sequences for peptide library **A**

**Spot	sequence	Spot	sequence**

```
-----------------------	-----------------------
A-1	AHGSGAWLLPV	A-51	NNRLPSSASAL
A-2	VHGSGAWLLPV	A-52	QNRLPSSASAL
A-3	IHGSGAWLLPV	A-53	DNRLPSSASAL
A-4	LHGSGAWLLPV	A-54	ENRLPSSASAL
A-5	MHGSGAWLLPV	A-55	RNRLPSSASAL
A-6	FHGSGAWLLPV	A-56	KNRLPSSASAL
A-7	YHGSGAWLLPV	A-57	HNRLPSSASAL
A-8	WHGSGAWLLPV	A-58	CNRLPSSASAL
A-9	SHGSGAWLLPV	A-59	GNRLPSSASAL
A-10	THGSGAWLLPV	A-60	PNRLPSSASAL
A-11	NHGSGAWLLPV	A-61	RALGYLAGSAG
A-12	QHGSGAWLLPV	A-62	RVLGYLAGSAG
A-13	DHGSGAWLLPV	A-63	RILGYLAGSAG
A-14	EHGSGAWLLPV	A-64	RLLGYLAGSAG
A-15	RHGSGAWLLPV	A-65	RMLGYLAGSAG
A-16	KHGSGAWLLPV	A-66	RFLGYLAGSAG
A-17	HHGSGAWLLPV	A-67	RYLGYLAGSAG
A-18	CHGSGAWLLPV	A-68	RWLGYLAGSAG
A-19	GHGSGAWLLPV	A-69	RSLGYLAGSAG
A-20	PHGSGAWLLPV	A-70	RTLGYLAGSAG
A-21	AYLGYLAGSAG	A-71	RNLGYLAGSAG
A-22	VYLGYLAGSAG	A-72	RQLGYLAGSAG
A-23	IYLGYLAGSAG	A-73	RDLGYLAGSAG
A-24	LYLGYLAGSAG	A-74	RELGYLAGSAG
A-25	MYLGYLAGSAG	A-75	RRLGYLAGSAG
A-26	FYLGYLAGSAG	A-76	RKLGYLAGSAG
A-27	YYLGYLAGSAG	A-77	RHLGYLAGSAG
A-28	WYLGYLAGSAG	A-78	RCLGYLAGSAG
A-29	SYLGYLAGSAG	A-79	RGLGYLAGSAG
A-30	TYLGYLAGSAG	A-80	RPLGYLAGSAG
A-31	NYLGYLAGSAG	A-81	RAGSGAWLLPV
A-32	QYLGYLAGSAG	A-82	RVGSGAWLLPV
A-33	DYLGYLAGSAG	A-83	RIGSGAWLLPV
A-34	EYLGYLAGSAG	A-84	RLGSGAWLLPV
A-35	RYLGYLAGSAG	A-85	RMGSGAWLLPV
A-36	KYLGYLAGSAG	A-86	RFGSGAWLLPV
A-37	HYLGYLAGSAG	A-87	RYGSGAWLLPV
A-38	CYLGYLAGSAG	A-88	RWGSGAWLLPV
A-39	GYLGYLAGSAG	A-89	RSGSGAWLLPV
A-40	PYLGYLAGSAG	A-90	RTGSGAWLLPV
A-41	ANRLPSSASAL	A-91	RNGSGAWLLPV
A-42	VNRLPSSASAL	A-92	RQGSGAWLLPV
A-43	INRLPSSASAL	A-93	RDGSGAWLLPV
A-44	LNRLPSSASAL	A-94	REGSGAWLLPV
A-45	MNRLPSSASAL	A-95	RRGSGAWLLPV
A-46	FNRLPSSASAL	A-96	RKGSGAWLLPV
A-47	YNRLPSSASAL	A-97	RHGSGAWLLPV
A-48	WNRLPSSASAL	A-98	RCGSGAWLLPV
A-49	SNRLPSSASAL	A-99	RGGSGAWLLPV
```

A-50	TNRLPSSASAL	A-100	RPGSGAWLLPV

**SUPPLEMENTARY TABLE 4** Spot sequences for peptide library **B**

**Spot	sequence	Spot	sequence**

```
-----------------------	-----------------------
B-1	MAGSGAWLLPV	B-51	NYLGYLAGSAG
B-2	MVGSGAWLLPV	B-52	QYLGYLAGSAG
B-3	MIGSGAWLLPV	B-53	DYLGYLAGSAG
B-4	MLGSGAWLLPV	B-54	EYLGYLAGSAG
B-5	MMGSGAWLLPV	B-55	RYLGYLAGSAG
B-6	MFGSGAWLLPV	B-56	KYLGYLAGSAG
B-7	MYGSGAWLLPV	B-57	HYLGYLAGSAG
B-8	MWGSGAWLLPV	B-58	CYLGYLAGSAG
B-9	MSGSGAWLLPV	B-59	GYLGYLAGSAG
B-10	MTGSGAWLLPV	B-60	PYLGYLAGSAG
B-11	MNGSGAWLLPV	B-61	TANPPSAQIKP
B-12	MQGSGAWLLPV	B-62	TVNPPSAQIKP
B-13	MDGSGAWLLPV	B-63	TINPPSAQIKP
B-14	MEGSGAWLLPV	B-64	TLNPPSAQIKP
B-15	MRGSGAWLLPV	B-65	TMNPPSAQIKP
B-16	MKGSGAWLLPV	B-66	TFNPPSAQIKP
B-17	MHGSGAWLLPV	B-67	TYNPPSAQIKP
B-18	MCGSGAWLLPV	B-68	TWNPPSAQIKP
B-19	MGGSGAWLLPV	B-69	TSNPPSAQIKP
B-20	MPGSGAWLLPV	B-70	TTNPPSAQIKP
B-21	MARLPSSASAL	B-71	TNNPPSAQIKP
B-22	MVRLPSSASAL	B-72	TQNPPSAQIKP
B-23	MIRLPSSASAL	B-73	TDNPPSAQIKP
B-24	MLRLPSSASAL	B-74	TENPPSAQIKP
B-25	MMRLPSSASAL	B-75	TRNPPSAQIKP
B-26	MFRLPSSASAL	B-76	TKNPPSAQIKP
B-27	MYRLPSSASAL	B-77	THNPPSAQIKP
B-28	MWRLPSSASAL	B-78	TCNPPSAQIKP
B-29	MSRLPSSASAL	B-79	TGNPPSAQIKP
B-30	MTRLPSSASAL	B-80	TPNPPSAQIKP
B-31	MNRLPSSASAL	B-81	TAGSGAWLLPV
B-32	MQRLPSSASAL	B-82	TVGSGAWLLPV
B-33	MDRLPSSASAL	B-83	TIGSGAWLLPV
B-34	MERLPSSASAL	B-84	TLGSGAWLLPV
B-35	MRRLPSSASAL	B-85	TMGSGAWLLPV
B-36	MKRLPSSASAL	B-86	TFGSGAWLLPV
B-37	MHRLPSSASAL	B-87	TYGSGAWLLPV
B-38	MCRLPSSASAL	B-88	TWGSGAWLLPV
B-39	MGRLPSSASAL	B-89	TSGSGAWLLPV
B-40	MPRLPSSASAL	B-90	TTGSGAWLLPV
B-41	AYLGYLAGSAG	B-91	TNGSGAWLLPV
B-42	VYLGYLAGSAG	B-92	TQGSGAWLLPV
B-43	IYLGYLAGSAG	B-93	TDGSGAWLLPV
B-44	LYLGYLAGSAG	B-94	TEGSGAWLLPV
B-45	MYLGYLAGSAG	B-95	TRGSGAWLLPV
B-46	FYLGYLAGSAG	B-96	TKGSGAWLLPV
B-47	YYLGYLAGSAG	B-97	THGSGAWLLPV
B-48	WYLGYLAGSAG	B-98	TCGSGAWLLPV
B-49	SYLGYLAGSAG	B-99	TGGSGAWLLPV
B-50	TYLGYLAGSAG	B-100	TPGSGAWLLPV
```

**SUPPLEMENTARY TABLE 5** Putative MetAP and LFTR substrates bearing Nt sequence, (MTN)

| **Protein name (accession number)** | **Function** | **N-terminal Sequence (ORF)** | **Met cleavage** ^§^ **(Yes / No / n.d.)** | **Subcellular location** |
| --- | --- | --- | --- | --- |
| BarA (P0AEC5) | Two-component regulatory system UvrY/BarA involved in the regulation of carbon metabolism. | MTNYSLRARM | n.d. | Cytoplasmic |
| GalF (P0AAB6) | UTP--glucose-1-phosphate uridylyltransferase | MTNLKAVIPV | Yes | Cytoplasmic |
| PtkA (P69828) | Galactitol-specific phosphotransferase enzyme IIA component (GatA). | MTNLFVRSGI | Yes (partial) | Cytoplasmic |
| LpxL (P0ACV0) | Lipid A biosynthesis lauroyltransferase (HtrB) | MTNLPKFSTA | n.d. | Cytoplasmic |
| PriB (P07013) | Primosomal replication protein N | MTNRLVLSGT | Yes | Cytoplasmic |
| YbdL (P77806) | Methionine aminotransferase | MTNNPLIPQS | Yes | Cytoplasmic |
| CsdE (P0AGF2) | Sulfur acceptor protein CsdE (ygdK) | MTNPQFAGHP | Yes | Cytoplasmic |
| YgeI (Q46789) | Uncharacterized protein YgeI | MTNPIGINNL | n.d. | n.d. |
| TsgA (P60778) | Protein TsgA (YhfC) | MTNSNRIKLT | n.d. | Multi-pass IM protein (Nt, cytoplasmic) |
| EngB (P0A6P7) | Probable GTP-binding protein EngB (YihA) | MTNLNYQQTH | Yes (partial) | Cytoplasmic |
| YjiP (P0DP21) | Putative inactive recombination-promoting nuclease-like protein (YjiP) | MTNFTTSTPH | n.d. | n.d. |
| Gre1 (P32674) | Probable dehydratase (PflD) | MTNRISRLKT | n.d. | Cytoplasmic |

^§^ (Bienvenut et al., 2015)

### REFERENCES

Bienvenut, W.V., Giglione, C., and Meinnel, T. (2015). Proteome-wide analysis of the amino terminal status of Escherichia coli proteins at the steady-state and upon deformylation inhibition. *Proteomics* 15**,** 2503-2518.

Daigle, D.M., and Brown, E.D. (2004). Studies of the interaction of Escherichia coli YjeQ with the ribosome in vitro. *J Bacteriol* 186**,** 1381-1387.

Link, A.J., Robison, K., and Church, G.M. (1997). Comparing the predicted and observed properties of proteins encoded in the genome of Escherichia coli K-12. *Electrophoresis* 18**,** 1259-1313.
